# ECHO: an Application for Detection and Analysis of Oscillators Identifies Metabolic Regulation on Genome-Wide Circadian Output

**DOI:** 10.1101/690941

**Authors:** Hannah De los Santos, Emily J. Collins, Catherine Mann, April W. Sagan, Meaghan S. Jankowski, Kristin P. Bennett, Jennifer M. Hurley

## Abstract

**Motivation:** Time courses utilizing genome scale data are a common approach to identifying the biological pathways that are controlled by the circadian clock, an important regulator of organismal fitness. However, the methods used to detect circadian oscillations in these datasets are not able to accommodate changes in the amplitude of the oscillations over time, leading to an underestimation of the impact of the clock on biological systems.

**Results:** We have created a program to efficaciously identify oscillations in large-scale datasets, called the Extended Circadian Harmonic Oscillator application, or ECHO. ECHO utilizes an extended solution of the fixed amplitude mass-spring oscillator that incorporates the amplitude change coefficient. Employing synthetic datasets, we determined that ECHO outperforms existing methods in detecting rhythms with decreasing oscillation amplitudes and recovering phase shift. Rhythms with changing amplitudes identified from published biological datasets revealed distinct functions from those oscillations that were harmonic, suggesting purposeful biologic regulation to create this subtype of circadian rhythms.

**Availability:** ECHO’s full interface is available at https://github.com/delosh653/ECHO. An R package for this functionality, echo.find, can be downloaded at https://CRAN.R-project.org/package=echo.find.

**Supplementary information:** Supplementary data are available

## 1 Introduction

Circadian rhythms are 24-hour oscillations that allow for the anticipation of the day/night cycle. They confer an evolutionary advantage by enabling an organism to optimize the timing of their cellular physiology such that biological processes occur at advantageous times (Dunlap (1999)). Daily oscillations occur in many processes, including metabolic regulation, immune function, and sleep (Decoursey *et al.* (1997); Klarsfeld and Rouyer (1998); Levi *et al.* (2010); Ouyang *et al.* (2009)). Conversely, chronic disruption of circadian rhythms is strongly associated with an increased risk of disease development, e.g. cancer, diabetes, and cardiovascular disease (Evans and Davidson (2013)). Circadian rhythms are generated at the molecular level by a highly conserved circadian “clock” comprising a transcription-translation negative feedback loop that cycles once every 24 hours (Hurley *et al.* (2016); Partch *et al.* (2014)). While it is known that there is widespread regulation originating from the components of the clock at both the transcriptional and translational levels, termed the clocks “output”, the field is only beginning to appreciate the complexity of the cellular circadian regulatory network (e.g.Hurley *et al.* (2014); Robles *et al.* (2014); Wang *et al.* (2017); Mure *et al.* (2018)).

In higher eukaryotes, the circadian clock exists in each cell and these independent clocks are synchronized by a master regulator in the brain, e.g. the Suprachiasmatic Nucleus (SCN) in mammals (Antle and Silver (2005); Ralph *et al.* (1990); Kowalska and Brown (2007)). While the master regulators maintain a robust circadian rhythm regardless of their environment, rhythms in peripheral tissues have been suggested to damp, or decrease in amplitude over time, without persistent entrainment cues (Aton *et al.* (2005); Yamazaki *et al.* (2000)). Thus, under LD or similar entrainment conditions, clock and clock-regulated genes are observed to be harmonic, whereas without these cues many genes have been observed to damp in their oscillation over time. A debate as to the cause of this damping is still underway, suggesting either global circadian damping or the desynchronization of individual cell rhythms as the cause, with recent models being unable to reject either hypothesis (Westermark *et al.* (2009)). Regardless of the cause, models of the clock have predicted that damping is an essential part of proper circadian time-keeping and reports have shown that damping plays a role in proper biological timing, including photoperiodicity, temperature compensation, and entrainment (Beer *et al.* (2017); Horikawa *et al.* (2005); Tseng *et al.* (2012)). However, the impact of circadian damping has not been explored on the output of the clock, presumably due to a lack of computational tools to examine damping in high-throughput datasets (Loros *et al.* (2007)).

To investigate clock output (referred to here as clock-controlled elements or CCEs) researchers conduct comprehensive omics analyses over circadian time (e.g. Hurley *et al.* (2014); Jang *et al.* (2015); Menet *et al.* (2012); Robles *et al.* (2014); Rodriguez *et al.* (2012); Hurley *et al.* (2018)). To enable analysis of these datasets, researchers take advantage of the idea that CCEs exhibit oscillatory patterns that follow sinusoidal curves and categorize data as rhythmic based on the similarity between an oscillation and reference curves, such as cosine curves (Hughes *et al.* (2010); Hutchison *et al.* (2015); Wu *et al.* (2016)). However, most CCE detection methods cannot take into account damping. Moreover, CCE detection methods that can account for damping are not effective on a high-throughput scale, as these approaches generally work directly with the original differential equation, creating a computationally expensive method (Westermark *et al.* (2009); Eser *et al.* (2014)). Therefore, the current circadian detection algorithms likely miss many damped CCEs.

To better elucidate CCEs, we created the Extended Circadian Harmonic Oscillator (ECHO) application (app), a publicly-available, easy-to-use interface that rapidly detects rhythms with and without a change in amplitude in large scale datasets. ECHO uses an optimization approach that fits the solution to the differential equation corresponding to an underlying negative feedback loop with external influences to omics scale data, thus extending the simple harmonic equation to identify the degree of amplitude change. ECHO detects not only the change in amplitude of oscillating genes, but it is more robust to noise than current circadian detection algorithms. Applying ECHO to previously published large scale transcriptomic and proteomic datasets, we detected CCEs that fell into damped, harmonic, and even forced (oscillations that increase in amplitude over circadian time) groups. CCEs that were classified as damped or forced fell into distinct Gene Ontological categories, even across several different species. Moreover, we found the environmental conditions of sampling impacted whether genes are damped or forced. Finally, we uncovered potential transcriptional connections stemming from the clock that could lead to the differential regulation of clock output, suggesting that damping is primarily a directed process rather than a by-product of desynchronization.

## 2 Methods

State-of-the-art methods for the identification of circadian rhythms in large scale datasets utilize fixed amplitude models (Hughes *et al.* (2010); Hutchison *et al.* (2015); Wu *et al.* (2016)). These methods are encompassed by the solution to the simple harmonic oscillator differential equation, a fixed amplitude model:

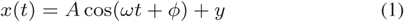

where *x*(*t*) is the resulting change in amplitude at time *t, A* is the initial amplitude, *ω* is the frequency of oscillation, *ϕ* is the phase shift, and *y* is the equilibrium value. This solution to the harmonic oscillator differential equation (1) corresponds to an underlying negative feedback loop (Pigolotti *et al.* (2007)), with a set of activators and repressors corresponding to the rise and fall of an oscillation’s trends. Though this equation can capture the oscillations in large-scale circadian datasets, it cannot account for outside forces that can change the amplitude of a negative feedback loop over time. However, it is important to note this system can be (and in reality, almost always is) influenced by outside biological forces, such as experimental conditions, that can decrease (damp) or increase (force) the amplitude of the oscillation it is describing over time. For example, when organisms are kept in “constant conditions” (e.g. darkness or without a media change that can lead to nutrient depletion), these environmental factors have the potential to force the amplitude of the oscillation to change (Aton *et al.* (2005); Yamazaki *et al.* (2000)). To account for these more realistic biological conditions in the identification of oscillations in large scale datasets, we used the extended harmonic oscillator equation:

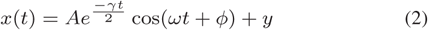

where *γ* is the amplitude change (AC) coefficient, and all other parameters retain their meaning from (1). Of the utmost importance in this equation is the inclusion of the amplitude change coefficient, which quantifies the amount of change in amplitude within the system. A positive value of *γ* indicates damping, or decreasing amplitude over time, and a negative value of *γ* indicates forcing or increasing amplitude over time. This equation reduces to (1) in the ideal case, as a truly harmonic oscillator has a *γ* value of 0. By utilizing (2), changes in amplitude can be accounted for, leading to the more complete identification of circadian rhythms. In addition, by using (2) we can capture the magnitude of *γ* to categorize each oscillation as damped, forced or harmonic.

### 2.0.1 Fitting Data with the Extended Harmonic Oscillator Equation

Building off the approach proposed in (De los Santos *et al.* (2017)), to find the parameter values in (2), we use the method of nonlinear least squares. Given experimental data **(t**,**x(t))** = (*t*_1_, *x*(*t*_1_)), (*t*_2_, *x*(*t*_2_)), *…*, (*t*_*n*_, *x*(*t*_*n*_)) and parameters *β* = (*A, γ, ω, ϕ, y*), this method minimizes the squared difference between experimental and fitted data in the following manner:

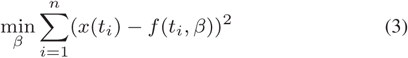

where *n* is the total number of data points and *f* (*t*_*i*_, *β*) refers to (2). We find a local solution to this minimization problem using the package `minpack.lm` in `R`, which uses the Levenberg-Marquadt algorithm for nonlinear least squares (Elzhov *et al.* (2016)). Confidence intervals for all parameters can be computed by bootstrapping, as computed by the package `nlstools` (Baty *et al.* (2015)).

As nonlinear least squares is a nonconvex problem, choosing the correct starting values is important. We thus choose our starting points for each of the parameters (denoted with subscript 0) with heuristics based on the experimental data (Supplemental Section 1).

### 2.0.2 Analyzing Datasets with Multiple Replicates

The equation given above (3) for nonlinear least squares assumes that for *β* to be an unbiased estimator, the variance at each time point must be equal (Strutz (2010)). In the case where there is a single replicate, this is vacuously true, as each replicate with a single time point must have zero variance. However, if a given dataset has multiple measurements at each time point, one cannot assume that variances are equal. To compensate for this discrepancy and obtain an unbiased *β*, we add weights at each time point equal to the variance at each time point (Strutz (2010)). Thus, (3) becomes:

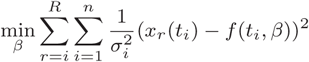

where *R* is the total number of replicates, 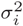 is the estimated variance at each time point, and *x*_*r*_(*t*_*i*_) represents the expression value of replicate *r* at time point *i*, and all other variables retain their meaning from (3). From first principles, we surmise that we should “trust” time points more if their values are closer together (lower variance) rather than far apart (higher variance).

### 2.0.3 Estimating Goodness of Fit

Once the parameters are obtained, we determine goodness of fit by computing the P-value using Kendall’s tau correlation coefficient, which measures the concordance between two series of data. In this case, we compare the fitted data values y = (*y*_1_, …, *y*_*n*_) to the experimental data values x = (*x*_1_, …, *x*_*n*_), with *n* corresponding to the total number of data points in the series. Then Kendall’s *τ* is measured as follows (Hutchison *et al.* (2015)):

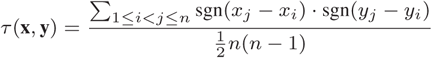

where sgn(*x*) is 1 if *x* is positive and −1 if *x* is negative. The numerator corresponds to the number of pairs that vary concordantly minus the number that vary discordantly. The denominator corresponds to the total amount of comparisons, such that *τ* is normalized to be between −1 and 1. P-values are calculated using the exact Kendall’s tau statistic. P-values are then adjusted, either using the Benjamini-Hochberg or Benjamini-Yekutieli criteria to account for multiple hypothesis testing, and evaluated for significance using a 0.05 cutoff.

### 2.0.4 Identifying the Harmonics of Oscillating Elements

Once a fit is determined to be significant, the rhythm is categorized based on its value of the amplitude change coefficient (*γ*). We determined empirically that if 0.15 ≤ *γ* ≤ 0.03, the rhythm should be categorized as as damped; if −0.15 ≤ *γ* ≤ −0.03 the rhythm should be categorized as as forced; if −0.03 ≤ *γ* ≤ 0.03, the rhythm should be categorized as as harmonic, though we feel that this determination may be dependent upon the noise in the data (see further discussion in Supplemental Section 3, Fig. 1A-C, 2A). Should *γ* be less than −0.15 or greater than 0.15 for a specific rhythm, the gene is considered to be overexpressed or repressed, respectively (Supplemental Fig.1D and E, 2B and C). Rhythms with these magnitudes of *γ* are not considered to be circadian, as their expression patterns more accurately represent a sharp increase or decrease in the amplitude with a small amount of noise rather than an oscillation over time (Supplemental Section 3).

**Fig. 1.**
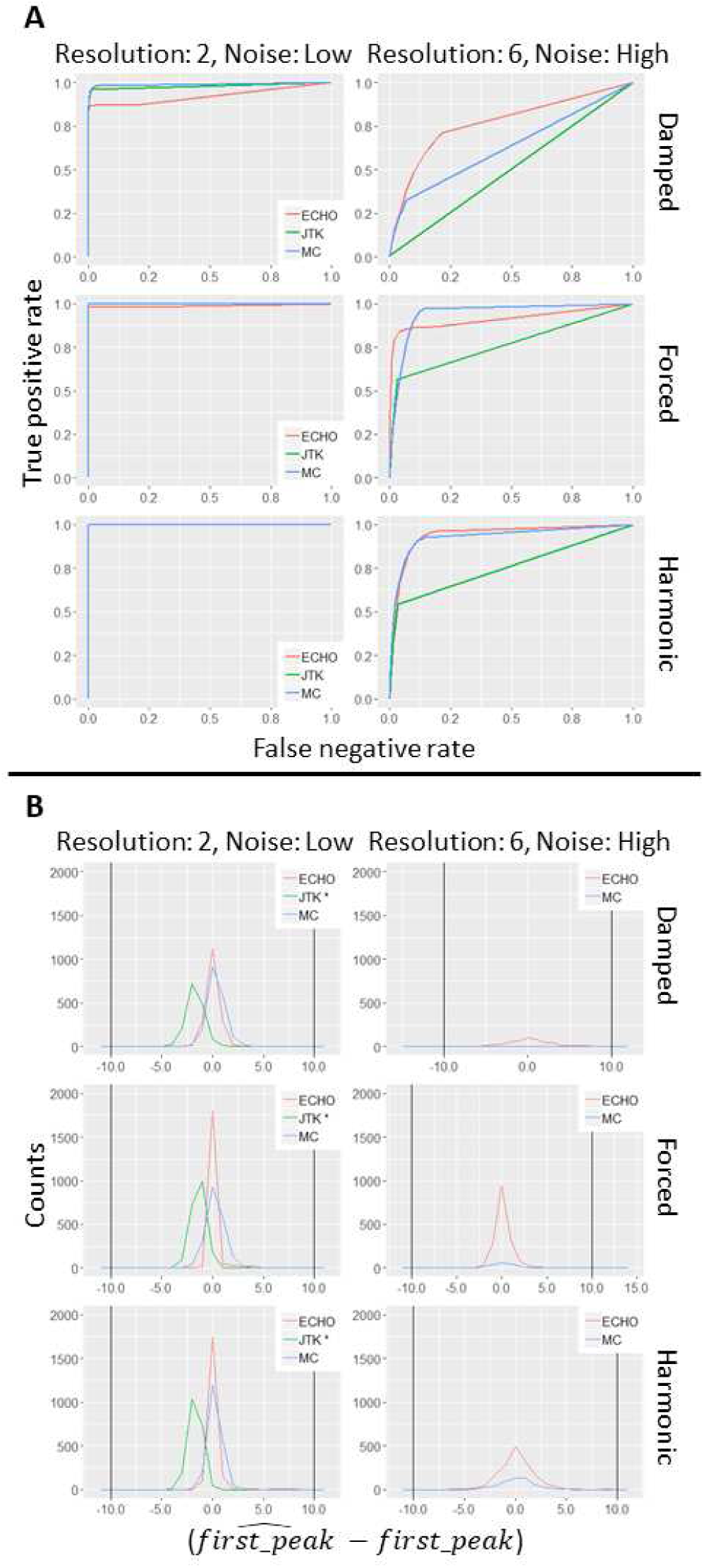
ROC curves demonstrate that ECHO robustly detects oscillations and precisely defines period. A. ROC curves for ECHO, JTK, and MetaCycle when analyzing expression patterns with low noise and 2-hour resolution or high noise and 6-hour resolution with amplitude change coefficients which fall into the damped, forced, and harmonic ranges, respectively, in a dataset with a 1:4 circadian to non-circadian ratio. B. Frequency polygon plots for ECHO, JTK, and MetaCycle when analyzing phase in data with low noise and 2-hour resolution or high noise and 6-hour resolution with amplitude change coefficients which fall into the damped, forced, and harmonic ranges, respectively, in a dataset with a 1:4 circadian to non-circadian ratio. ECHO = the ECHO method. JTK = the JTK_CYCLE method. MC = the MetaCycle method.

### 2.1 The Extended Circadian Harmonic Oscillator (ECHO) Application

We created an ECHO interface which allows researchers to both identify rhythms and visualize their results (Supplemental Section 4.1, Supplemental Fig. 3), available on GitHub^1^ and as an R package through CRAN (De los Santos *et al.* (2018)). The ECHO application allows for the use of a variety of preprocessing techniques. However, preprocessing, as with any editing of values, will likely change the estimated parameter values (Supplemental Section 4.2). ECHO’s interface allows for searches for rhythms of any period length up to the total length of the time course, a “free run”. (Supplemental Fig. 4A and B). Free runs also prevent suboptimal rhythms from being detected at a specified lower or higher period bound, when their optimal periods fit outside of the user specified range (Supplemental Section 4.3, Fig. 4C and D). It should be noted that the ECHO method works with any numerical data, including standard genome-wide data analysis techniques, such as microarray and RNA-seq, as well as longer time courses, such as real-time luciferase assays.

**Fig. 2.**
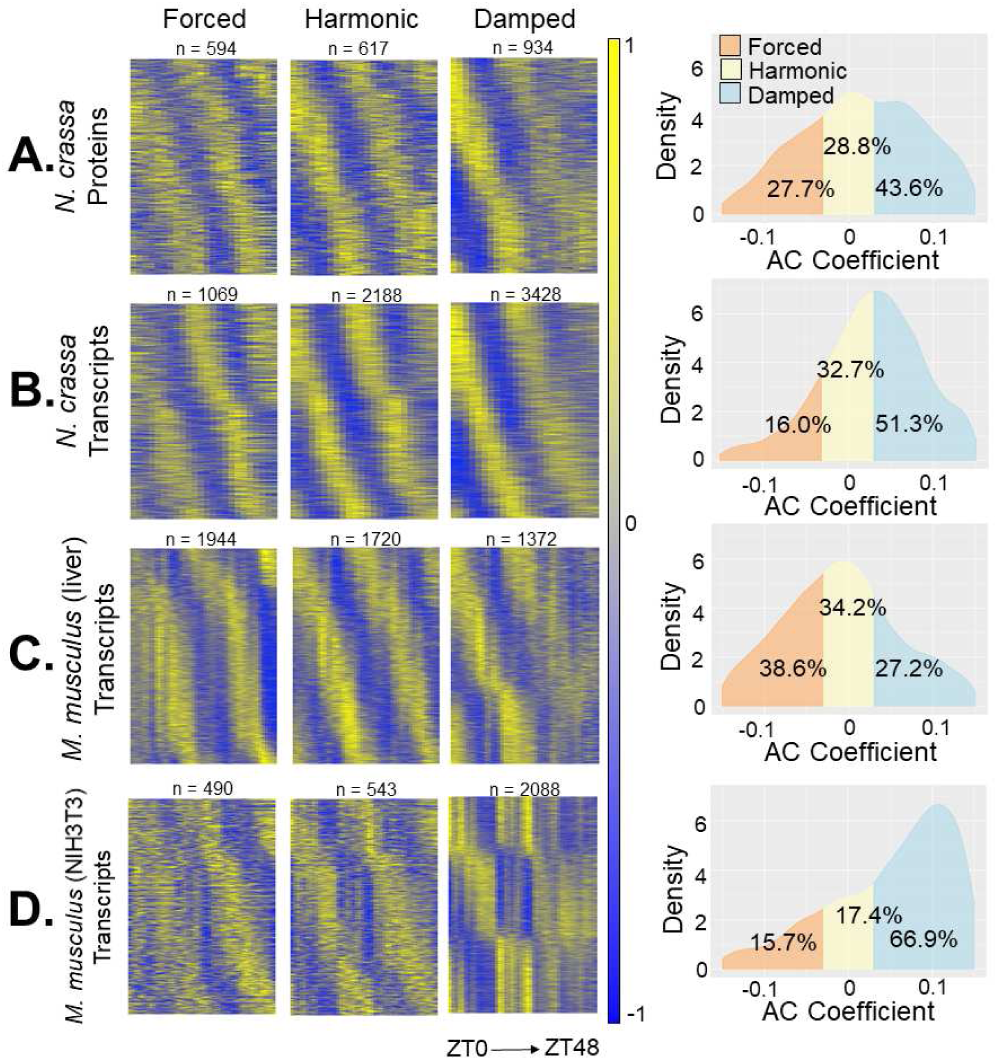
The ratio of harmonic to damped to forced CCEs depends on sampling conditions. Heatmaps and AC coefficient density graphs of the CCEs determined by ECHO to be circadian in A. the N. crassa proteome (Hurley et al. (2018)), B. the N. crassa transcriptome (Hurley et al. (2014)), C. the M. musculus liver transcriptome (Hughes et al. (2009)), and D. the M. musculus NIH3T3 transcriptome (Hughes et al. (2009)). For each dataset, the heat maps show mean-centered normalized expression values at a given time point for the transcripts that fall into the AC coefficient categories damped, forced, or harmonic, and are sorted vertically by phase.

**Fig. 3.**
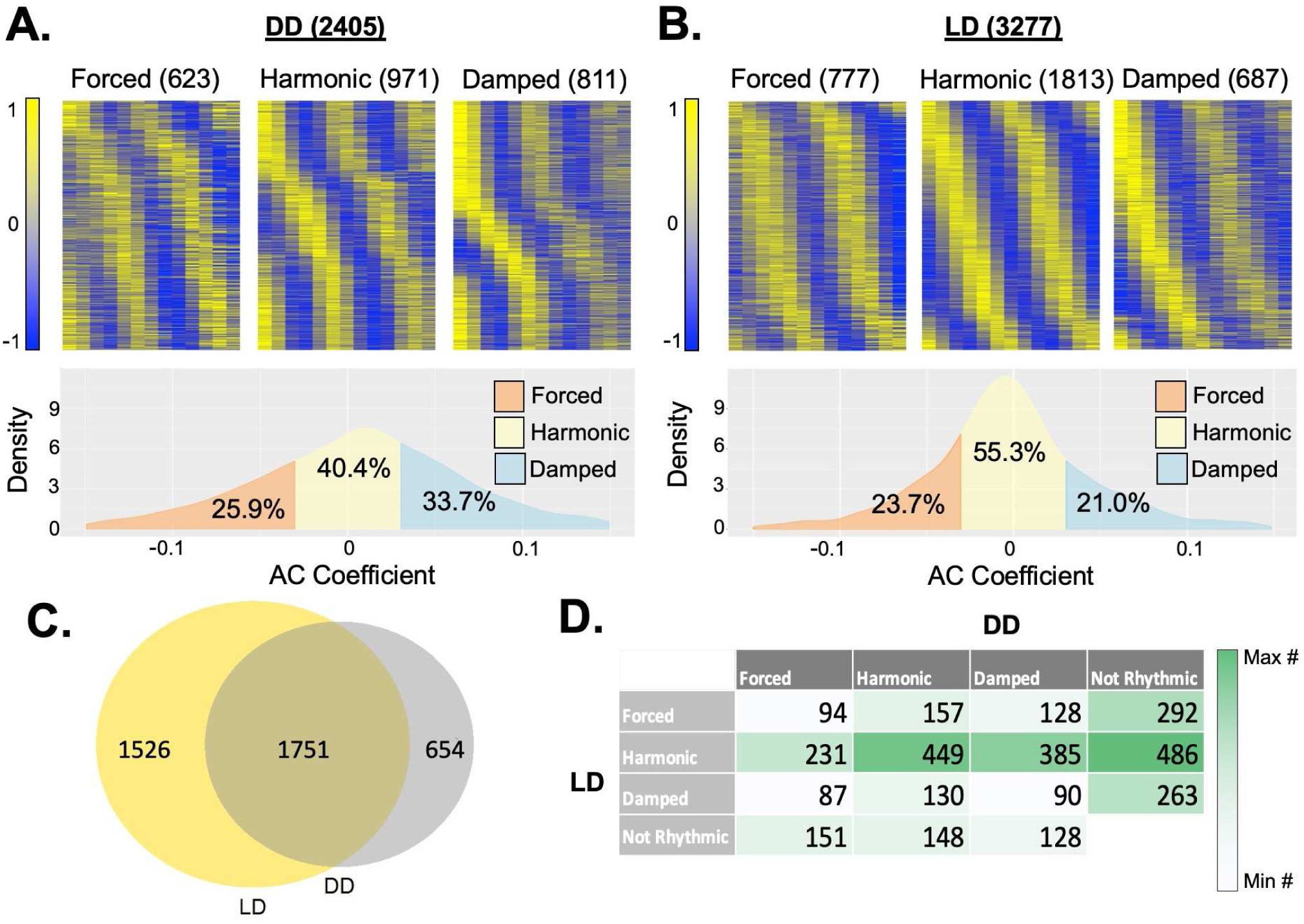
The ratio of damped, forced, and harmonic CCEs varies depending on lighting schemes in Anopheles gambiae. A. and B. Heat maps and AC coefficient density graphs of the transcripts defined as damped, forced, or harmonic by ECHO in Anopheles gambiae (mosquito) heads gathered in either complete darkness (DD) (A) or 12:12 Light/Dark (LD) (B)(Rund et al. (2011)). C. A Venn diagram describing the overlap of transcripts found to be rhythmic in Anopheles in DD and LD. D. A confusion matrix comparing the best fit models (damped, forced, and harmonic) for CCEs in Anopheles identified as circadian by ECHO in either DD or LD conditions.

**Fig. 4.**
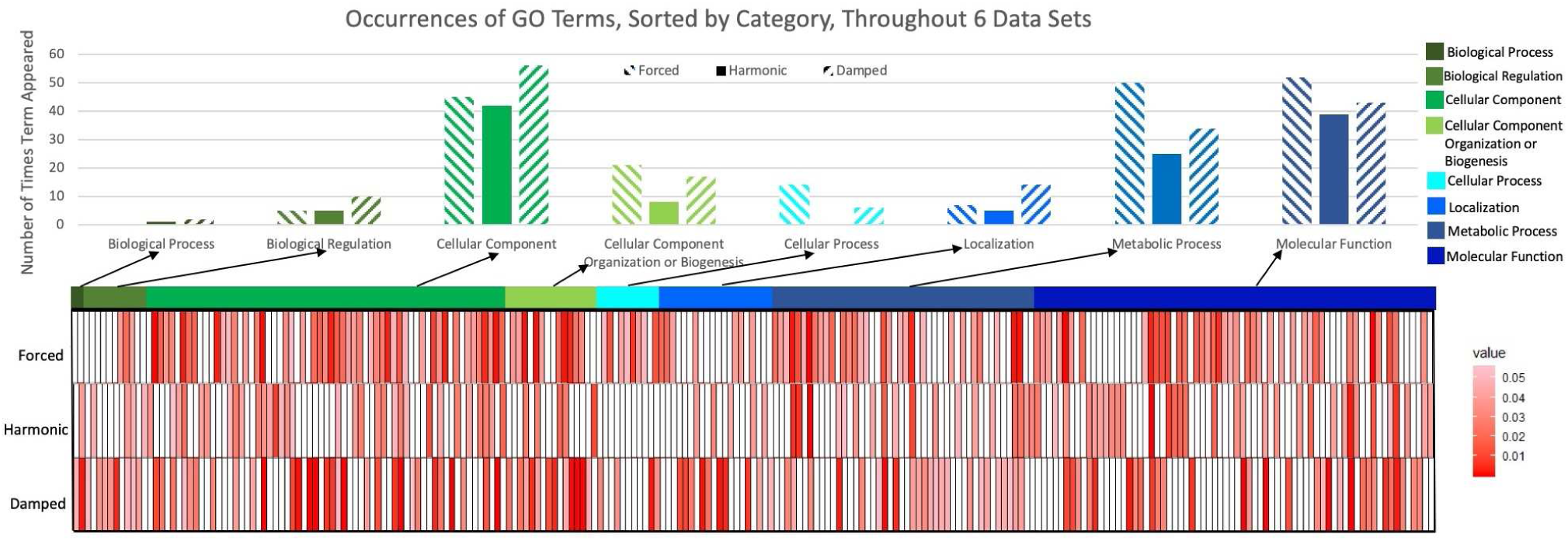
Gene Ontological analysis suggests specific biological functions for damped, forced, and harmonic CCEs. All transcriptomic data (microarray & RNA-seq derived) were included in a meta-analysis of Gene Ontologies. All significant (FDR adjusted *P* -*value* < 0.1) ontology subcategories were grouped by parent hierarchical term and the total number of CCEs in the corresponding categories are shown above in bar graphs. Heat maps show the enrichment P-value for the subcategories from a given parent ontology in A. Ontological subcategories are classified by their AC coefficient group, with exact false discovery rates given in Supplemental Data: Table 1.

## 3 Results

### 3.1 ECHO Exceeds Current Methodologies in Rhythm and Phase Detection

To evaluate the capabilities of ECHO as compared to existing approaches (Hughes *et al.* (2010); Wu *et al.* (2016)), we applied ECHO, JTK, and MetaCycle to synthetically generated datasets with circadian and non-circadian data (Supplemental Section 5.1) and calculated the area under the receiver operating characteristic (ROC) curve (AUC) for all three programs. We utilized two basic synthetic dataset structures, one that consisted of a realistic circadian to noncircadian ratio and total number of genes (ratio of circadian:non-circadian of 1:4, 10,000 total expression patterns), and one that allowed us to examine the effects of multiple hypothesis testing (ratio of circadian:non-circadian of 1:1, 4,000 total expression patterns). Within each of these datasets, we further varied the noise levels (low, medium, and high), the time point resolution (2, 4, and 6 hours), and the type of circadian Amplitude Change (AC) categories (damped, forced, and harmonic) that were associated with each of the gene expression patterns.

In the dataset with a circadian to non-circadian ratio of 1:4, a damped AC coefficient, low level of noise, and high resolution sampling, each of the three programs performed well (Fig. 1A, Supplemental Table 1). However, as noise increased and resolution decreased to levels similar to what is seen in the literature, ECHO outperformed JTK and MetaCycle (Fig. 1A, Supplemental Table 1). To explore the effects of multiple hypothesis testing on ECHO, we examined the 1:1 circadian to non-circadian ratio dataset with ECHO. As in the 1:4 dataset, ECHO strongly outperformed the other two analysis methods (Supplemental Table 4).

In the case of the forced and harmonic AC coefficient datasets, at low noise and high resolution, JTK, Metacycle, and ECHO all maintained similar AUC scores in both the 1:4 and 1:1 datasets (Fig. 1A, Supplemental Tables 2, 3, 5, and 6). As the noise was increased and the resolution decreased, ECHO demonstrated similar and slightly higher AUC scores than Metacycle, and ECHO and Metacycle consistently outperformed JTK (Fig. 1A, Supplemental Tables 2, 3, 5, and 6). In the cases where ECHO did not have the highest AUC, it was numerically very similar to the highest, being no more than 0.08 from the AUC highest score (Supplemental Tables 1-6).

We tested the accuracy of each program to recover phase by computing the difference between the predicted phase shift from each program and the original generated phase shift for detected circadian genes with a BH-adjusted P-value cutoff of 0.05 (Supplemental Section 5.1). For our 1:4 ratio data, both ECHO and MetaCycle have distributions centered around 0 with a relatively low spread (Fig. 1B; Supplemental Figs. (5-7)A). This spread increases with noise and resolution, regardless of amplitude change coefficient category, though ECHO consistently recovers more circadian genes than MetaCycle. JTK, however, has a distribution centered between −1 and −2 with a low spread, indicating that JTK consistently underestimates phase. It also identifies far fewer circadian genes than ECHO and Metacycle, to the point that at high noises and resolutions, no synthetic circadian expression patterns are recovered by JTK. We noted that ECHO retains accuracy and continues to identify circadian genes at high noise and wider hour resolution for damped genes, and maintains this robustness for all amplitude change coefficient categories. In our 1:1 ratio data, ECHO, MetaCycle and JTK identify phase and gene numbers similar to the 1:4 ratio dataset; these observations are consistent with the observations from our ROC curves (Fig. 1B; Supplemental Fig. (5-7)B).

### 3.2 ECHO Analysis with Biological Data Reveals Extent and Sources of Damping and Forcing

To demonstrate ECHO’s increased efficiency and determine the extent of damping in circadian networks, we applied ECHO to publicly available 48-hour time course transcriptomic and proteomic datasets gathered from *Neurospora crassa* (Supplemental Section 5.2, 6.2, 6.3)(Hurley *et al.* (2014, 2018)). Of the 4,747 *Neurospora* proteins that were detected by tandem mass tag mass spectrometry, ECHO identified 2,146 CCEs, 935 (43.6%) of which were damped, 594 (27.7%) were forced, and 617 (28.8%) were harmonic (Fig. 2A). From the *Neurospora* transcriptome, ECHO identified 6,685 CCEs with a comparable proportion of harmonic CCEs (32.7%), a lower proportion of forced CCEs (16.0%), and a higher proportion of damped CCEs (51.3%)(Fig. 2B, Supplemental Fig. 8A) as compared to the proteome. Of note, multiple circadian clock genes were identified as damped (Supplemental Fig. 9, Supplemental Section 6.5). To examine if damping/forcing of genes at the transcriptional level is also reflected at the translational level, we created a confusion matrix of CCEs that were detected and modeled in both datasets. In addition to the damped transcripts that were not rhythmic (29.3%), or damped/repressed at protein level as expected (38.6%), a comparable proportion were also found to be harmonic, forced or overexpressed (31.9%) despite damping at the transcriptional level (Supplemental Table 7). These differences between the proportion of damped and forced CCEs at the transcriptional and post-transcriptional levels suggested that there is extensive differential regulation of CCE expression by the circadian system to generate damping or forcing in a directed manner.

When we compared the ECHO analysis of the *Neurospora* transcriptome with a JTK analysis of the same transcriptomic data, we found that ECHO identified 882 CCEs in addition to the 7,441 CCEs identified by both ECHO and JTK, including those condsidered overexpressed or repressed by ECHO. Only 397 CCEs found by JTK were not encompassed by ECHO. Of the 882 CCEs missed by JTK, the majority, 62.2%, were determined by ECHO to be damped, 24.8% were forced and 12.9% harmonic, which is consistent with our synthetic data finding that ECHO has an improved capability to detect damped gene expression patterns that cannot be detected with existing methods, identifying many additional CCEs.

We hypothesized that the environmental conditions under which the samples were acquired may impact the AC of output genes, as the *Neurospora* time course was sampled in a depleting media condition. Therefore, we analyzed a high-resolution transcriptomic analysis of pooled liver samples taken *in vivo* from mice in complete darkness and compared the ratio of damped/forced/harmonic CCEs to an *in vitro* transcriptomic time course with NIH3T3 mouse fibroblast cells acquired with identical depth/duration of sampling (Hughes *et al.* (2009)). In the mouse liver dataset, a total of 20,662 liver transcripts were analyzed by ECHO and 5,036 were identified to be CCEs (Fig. 2C). The majority of CCEs were classified as forced (38.6%), 34.2% classified as harmonic, and only 27.2% classified as damped. In contrast, the NIH3T3 time course had more than double the proportion of damped CCEs (66.9%), and far smaller proportions of harmonic (17.4%) and forced (15.7%) CCEs (Fig. 2D). Though these are very different tissue types, the overall trends indeed suggest that nutrient sensing mechanisms are integrated with the clock to globally dampen rhythmic gene expression in nutrient limited conditions.

To probe additional environmental sources that may impact the damping/forcing ratio, and because we noted a strong correlation between the light response and damped CCEs, we investigated the influence of light entrainment cues on circadian output AC. We focused on two transcriptomic time courses done in *Anopheles gambiae* that differ only in their lighting schemes: either maintaining a program of 12 hrs of light followed by 12 hrs of dark (LD) or complete darkness (DD) throughout the time course (Rund *et al.* (2011)). We identified 2,405 CCEs from the DD dataset (Fig. 3A, Supplemental Fig. 8B) and 3.277 CCEs from the LD dataset (Fig. 3B), suggesting that over all, light cues increase the number of CCEs. As would have been expected if continued light entrainment prevents damped and forced rhythms, there was a lower proportion of harmonic CCEs (40.4% vs. 55.3%) and a higher proportion of damped CCEs (33.7% vs 21%) in the DD dataset compared to the LD dataset. In addition, the distribution of amplitude change coefficients for each dataset’s CCEs showed a wider spread in DD (1st quartile: - 0.032, 3rd quartile: 0.044) than in LD (1st quartile: −0.028, 3rd quartile: 0.023), demonstrating that AC is more pronounced in the absence of light entrainment cues (Fig. 3A, B).

To ensure that the increase in CCEs under LD conditions was not due to differences in microarray detection and/or number of transcripts analyzed by ECHO, we confirmed that there were similar numbers of transcripts that were unexpressed (i.e. detected in *<* 70% of time points) or with no deviation (having completely flat expression across the time course). We then investigated the overlap of CCEs between the two lighting conditions and found that the majority of CCEs are rhythmic in both conditions (1,751), though there were also some that were rhythmic only under LD conditions (1,526) or only under DD conditions (654) (Fig. 3C). To further parse the expression patterns observed for each CCE under one condition compared to the other, we created a confusion matrix for the 3,412 total unique CCEs detected and modeled with ECHO in both datasets (Fig. 3D). For the CCEs that were rhythmic in LD, the majority of damped, forced or harmonic CCEs were found to be not rhythmic under DD conditions, indicating that light responsive genes were highly prevalent in all three expression categories. As would be expected if desynchronization were a major factor in damping, many of the forced, damped and harmonic LD CCEs that do not lose rhythmicity, become damped under DD conditions. However, of the CCEs that were forced, harmonic or damped under LD conditions, the largest proportions (128, 449, 130, respectively) remained harmonic under DD conditions, suggesting that most circadian genes in fact do not become acutely damped or forced without synchronization cues and that intercellular desynchronization many not be responsible for circadian damping.

### 3.3 Meta-Analysis of Damped, Forced, and Harmonic Gene Ontologies Identifies Distinct Functions

To identify differences in the functional roles of the AC coefficient categories, we conducted a gene ontology enrichment meta-analysis of the CCEs identified in the four transcriptional datasets tested with ECHO (Supplemental section 5.3.1). The ontologies enriched in each AC category were found to be highly category-specific and suggestive of distinct functional roles for damping or forcing CCEs (Fig. 4). Distinct functional differences between the forced, damped, and harmonic CCEs became apparent in analyzing the enrichment of child-level ontology categories. Out of 247 total unique child ontology terms significant in at least one subset, the majority of terms (65%) were enriched in only one of the damped/forced/harmonic subsets, highlighting the unique specificity of gene functions present in each subset.

We examined the top twenty scoring ontologies unique to each expression pattern to further characterize the divergent biological roles of genes with damping, forcing, or harmonic expression (Supplemental Section 6.2). Most notable from the enriched damped CCEs (Supplemental Table 8) were several terms related to protein transport and localization within the cell (intracellular protein transport, localization, regulation of cellular component organization, macromolecule localization), kinase and transferase activity (kinase activity, phosphotransferase activity, transferring alkyl or aryl (other than methyl) groups, as well as transcription (transcription DNA templated, rRNA binding, RNA binding) and ribosome. Alternatively, the ontological categories unique to forced CCEs (Supplemental Table 9) pertained to catalysis of phosphate groups into free phosphate, such as in fatty acid activation (pyrophophatase activity, hydrolase activity, in phosphorus-containing anhydrides). Also uniquely forced were genes involved in RNA processing and translation (mRNA processing, translation, mRNA binding, etc.), and finally ubiquitin-mediated proteolysis (protein ubiquitination, proteasome-mediated ubiquitin-dependent protein catabolic process). Lastly, harmonic CCEs (Supplemental Table 10) were most notably related to macromolecule metabolism (macromolecule metabolic process, hydrolase activity acting on ester bonds, disulfide oxidoreductase activity), ribosome biogenesis (preribosome, rRNA processing, ribosome biogenesis), actin cytoskeleton maintenance (actin binding, cytoskeleton, actin cytoskeleton) and ion transmembrane gradient maintenance (proton transmembrane transporter activity, ion transmembrane transport). The distinctions between the functions of genes found in each category, in particular their unique metabolic functions, hint that environment/nutrient sensing underlies the regulation of circadian damping and forcing as a directed response to metabolize and preserve specific cellular functions efficiently in times of nutrient stress.

### 3.4 Enriched Regulatory Motifs Suggest Transcriptional Regulation Contributes to Damped and Forced Oscillations in *Neurospora crassa*

We hypothesized that transcription factor (TF) activity might explain the differential regulation of the forced, harmonic, or damped gene expression profiles. We therefore applied Discriminative Regular Expression Motif Elicitation (DREME) analysis (Bailey (2011)) to the promoter regions of the three sets of CCEs identified by ECHO from the *Neurospora crassa* transcriptomic dataset to determine if there were enriched motifs (Supplemental Section 5.3.2). DREME analysis detected many similar versions of four previously identified circadian motifs (Hurley *et al.* (2014)): STACASTA, GVCAGCCA, GRCGGGA, GCRCTAAC. Of note, the previous STACASTA motif that was enriched for cell cycle processes (Hurley *et al.* (2014)), matches our TACASTA motif that was identified in all three sets of our CCE promoters. When we compared this motif to a database of *Neurospora* transcription factor binding motifs Weirauch *et al.* (2014), we found that it matches a known binding motif for the transcription factor *sgr-21* (NCU06173), which is a “forkhead” type transcription factor that is involved in the cell cycle in other fungi (eg. *S. cerevisiae*, Pramila (2006)) and has a slower growth rate when knocked out in *Neurospora* (Carrillo *et al.* (2017)).

We also identified several novel motifs that only occurred in either the forced, harmonic, or damped gene sets. In the forced gene set, the unique GATAAG motif can be bound by two different transcription factors, *nit-2* (NCU09068) and *asd-4* (NCU15829), and their orthologs have been shown to co-regulate genes in response to nitrogen levels in *Aspergillus nidulans* (Fu and Marzluf (1990); Wong *et al.* (2009)). Meanwhile, the unique harmonic motif, CGGSCGG, could be bound by *rrg-2* (NCU02413), an oxidative stress response regulator that like it’s yeast homolog, Skn7, may be regulated by cAMP/PKA (Weirauch *et al.* (2014); Tian *et al.* (2011); Basenko *et al.* (2018); Hanlon *et al.* (2011); Fan *et al.* (2015); Pérez-Landero *et al.* (2015)). Finally, a motif that was enriched in the damped gene set only, CCCSKC, encodes a binding site for the transcription factor *cre-1* (NCU08807). CRE-1 is involved in carbon catabolite repression (Cupertino *et al.* (2015); Sun and Glass (2011); Ziv *et al.* (2008)), and ECHO classified *cre-1* as harmonic (BH adjusted P-value = 1.34e-07; Supplemental Fig. 10). Cupertino et al. (2015) identified binding motifs for CRE-1 within genes involved in glycogen metabolism, including *gsn* (NCU06687), *gdn* (NCU00743), *gnn* (NCU06698), *gbn* (NCU05429) (all down-regulated by CRE-1), and *gpn* (NCU07027) (up-regulated by CRE-1) (Cupertino *et al.* (2015)). Importantly, our ECHO analysis confirmed that known targets of CRE-1 repression were identified as either damped or repressed, (*gdn, gnn*, and *gbn*), while a known target of CRE-1 activation, *gpn*, was up-regulated and classified as forced (Supplemental Fig. 10). The gene expression for *gsn* which is predicted to be down-regulated was considered harmonic, but it is known to be further regulated by phosphorylation (Cupertino *et al.* (2015)) and also shows a slightly decreasing amplitude over time (AC coefficient = 0.0043).

## 4 Discussion and Conclusion

Guidelines outlined by the circadian community for the analysis of genome-scale experiments emphasizes the need for robust models to identify rhythmic expression patterns (Hughes *et al.* (2017)). By including the amplitude change coefficient (2), ECHO answers this call by providing an easy-to-use, enhanced approach to detect circadian patterns in large datasets, providing a freely-available point-and-click application with visualization and exploration tools rather than the standard code tutorial (Supplemental Fig. 3). This approach enhancement is verified by both ECHO’s superior recovery of circadian rhythms from synthetic data and previously published biological data (Fig. 1, 2, 3), its robustness to noise, resolution, and multiple hypothesis testing (Fig. 1), and its ability to provide the amplitude change coefficient (Supplemental Fig. 1).

Strikingly, ECHO’s extensive identification of damped and forced rhythms suggests a previously unforeseen impact of circadian damping. We also found the overall proportion of damped transcripts as compared to forced or harmonic transcripts was dependent on the environmental conditions in which the samples were collected; e.g. nutrient depletion increased damping (Fig. 2). This finding corresponded with the observation that metabolic regulation can have a strong impact on the circadian clock and suggests novel ways that the clock interacts with its environment (Eckel-Mahan and Sassone-Corsi (2013)). However, as the proteome revealed less bias towards damping at the protein level than observed in their corresponding transcripts, this demonstrates that a large proportion of CCEs that are damped at the transcriptional level are buffered at the translational level by currently unknown mechanisms, as has been previously predicted (Hurley *et al.* (2018)). The corresponding changes in mRNA degradation and translation rates for these genes would need to be further investigated in exploring whether this resistance to damping is derived from gene-specific or global mechanisms. ECHO would be a valuable tool in accomplishing such future work.

When we examine the roles of circadian entrainment cues, i.e. light, in the damping and forcing of output, we found that the lack of continued entrainment significantly impacted the downstream regulation of circadian output, further highlighting how the environmental state of the organism can impact its circadian output (Fig. 3) (Rund *et al.* (2011)). However, we found that many of the rhythmic genes in LD were more strongly influenced by light, as opposed to circadian regulation, as they were not rhythmic under DD conditions. Of those in LD that were harmonic, a large majority remained harmonic in DD, supporting that the loss of entrainment cues does not acutely result in global damping of CCEs via intercellular desynchronization. Further, when comparing all datasets analyzed with ECHO representing a variety of organisms, tissues, and 48-hour experimental designs (Table 1), we found that experiments done in a static artificial media have higher proportions of damped CCEs than experiments done under conditions where organisms had access to nutrients Ad Libitum regardless of their lighting conditions. This indicated that the loss of nutrients has a greater impact on the global proportion of damping of CCEs than a loss of light-based entrainment cues.

**Table 1.**
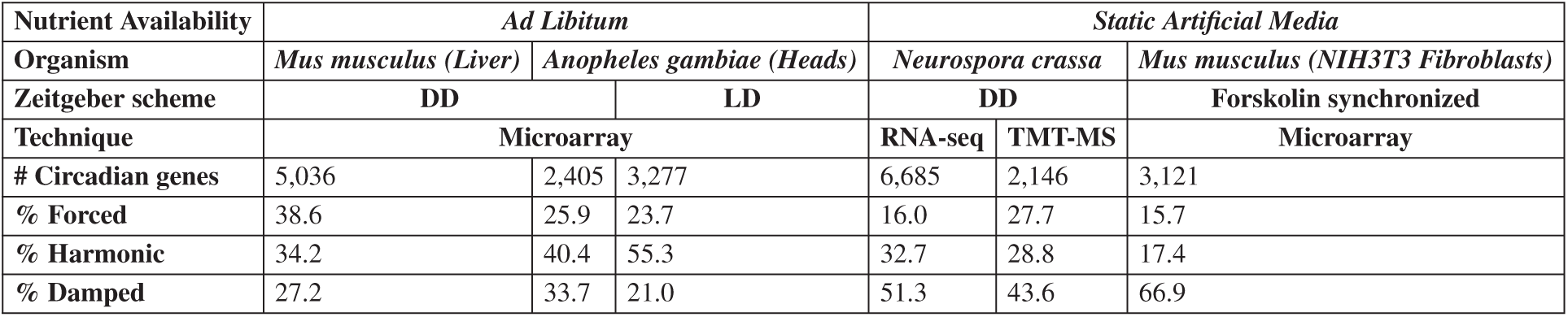
Proportions of Damped/Harmonic/Forced CCEs Vary By Experimental Conditions. Comparisons between relative number of CCEs found to be damped, forced, or harmonic, separated by experimental condition and organisms.

The observation of an impact on circadian output by the environment of the organism is bolstered by the distinct gene ontologies enriched in damped, forced, or harmonic transcripts (4). While the damping and forcing of presumably different cellular functions and metabolic pathways is indicative of a response to an environment with increasing stress from nutrient loss, of interest are the categories of genes that remain robustly harmonic. The harmonic preservation of genes involved in ribosome biogenesis, metabolism of macromolecules, ion transmembrane transport and actin cytoskeleton suggests a biological advantage to ensuring that these functions remain robust in all conditions. Alternatively, genes that dampen over time are involved in localization and bulk transcription and translation, suggesting that in order to preserve energy efficiently, it is advantageous to damp processes like localization and transcription/translation machinery when undergoing nutrient stress. Finally, genes that demonstrate forced expression over time involve the processing of mRNA, activation of fatty acids, and protein degradation, indicating that maintaining these processes ensures efficient, selective gene transcription and utilization of energy stored in fatty acids and proteins. Further, forced processes facilitate the robust degradation and recycling of protein and fatty acids, whereas harmonic processes serve to continue maintaining essential central macromolecular metabolic processes, ion gradients, cellular structure and pre-processing mRNA and creating ribosomes to ensure continued efficient expression of critical genes. Taken together, the distribution of these unique categories indicates that through damping/forcing/harmonic circadian regulation there is a built-in fine tuning of cellular processes to adapt advantageously to environmental changes.

Our DREME analysis on the *Neurospora* transcriptome suggests that *Neurospora* is using transcriptional programming to integrate environmental cues into circadian output (Hanlon *et al.* (2011)). Indeed, the activity of some of these transcription factors, such as RRG-2 and CRE-1, are regulated by conserved signaling pathways, suggesting again that forced or damped gene expression could be the outcome of sensed changes in the environment (e.g., Pérez-Landero *et al.* (2015)). As we know there is a connection between the clock and MAPK signaling, as well as some evidence for an interconnection between the clock and cAMP-PKA signaling, further studies using ECHO could help elucidate the important contributions of these and other signaling pathways to damped and forced circadian expressions (de Paula *et al.* (2008); O’Neill *et al.* (2008); Liu *et al.* (2015)).

In summary, we have demonstrated that ECHO analysis is a powerful tool to not only identify previously undetectable oscillations in genome-scale circadian datasets but also allows the user to gain insights into the architecture of the circadian system’s control of gene regulation and integration with environmental cues. Moreover, as ECHO is able to identify rhythms of almost any period (Supplemental Fig. 4), we predict that this easy-to-use application will allow a broad community to investigate the underlying principles of environmental signaling in biological rhythms.

## Acknowledgements

We would like to thank Carol Ringelberg for her guidance and expertise in pre-processing of microarray data. We would also like to thank Thomas Willemain for his statistical expertise, and Chendi Guan and Kristin Pratt for their assistance in developing and testing the ECHO application.

## Footnotes

1 https://github.com/delosh653/ECHO

