## Supplemental Text for "ECHO: an Application for Detection and Analysis of Oscillators Identifies Metabolic Regulation on Genome-Wide Circadian Output"

March 22, 2019

[1] Department of Computer Science, Rensselaer Polytechnic Institute, Troy, NY, U.S.A
[2] Institute of Data Exploration and Applications, Rensselaer Polytechnic Institute, Troy, NY, U.S.A
[3] Department of Biological Sciences, Rensselaer Polytechnic Institute, Troy, NY, U.S.A
[4] Department of Mathematical Sciences, Rensselaer Polytechnic Institute, Troy, NY, U.S.A
[5] Center for Biotechnology and Interdisciplinary Sciences, Rensselaer Polytechnic Institute, Troy, NY, U.S.A

#### 1 Starting Points for Nonlinear Least Squares

We developed a novel parameter initialization scheme for our nonconvex problem in (M.2). Our starting points for each of the parameters in (M.2)[1] (denoted with subscript 0) are based on heuristics in the averaged experimental data:

$$A_0 = \max(\mathbf{x(t)}) - \overline{\mathbf{x(t)}}$$

$$\gamma_0 = \begin{cases} \frac{1}{\sqrt{1+(\frac{2\pi}{\delta})^2}}, & \text{if \# of peaks } \geq 2\& \text{ peak}(1) > \text{ peak }(2) \\ \frac{-1}{\sqrt{1+(\frac{2\pi}{\delta})^2}}, & \text{if \# of peaks } \geq 2\& \text{ peak}(1) < \text{ peak }(2) \\ 0.01, & \text{if peaks } = 1 \\ 10^{-8}, & \text{otherwise} \end{cases} \tag{1}$$

$$w_0 = \frac{2\pi}{(\text{\# of time points})(\text{resolution})/(\text{\# of peaks})} \tag{2}$$

$$y_0 = \overline{\mathbf{x(t)}}$$

where # means number. In (2), resolution is defined as the difference between adjacent time points. $\phi_0$ is chosen by partitioning the possible phase shift range into 12 parts: $\frac{i\pi}{6}$, $i = 0, ..., 11$. Using the previously decided initial values, $\phi_0$ is then chosen based on the $i$ that minimizes the sum of the absolute value of the difference between the experimental data and the proposed initial fit, calculated using (M.2), at time points 2 and 3.

(1) and (2) include the vector of peaks within the experimental data and the logarithmic decrement. A data point is determined to be a peak if it is the maximum value of a set including itself and a certain number of surrounding data points. The size of this set is dependent on the resolution of the data. For 5 minute resolution or less, this set is size 102, for (5,10] minute resolution, size 45; for (10,15] minute resolution, size 26; (15,30] minute resolution, size 11, for (.5,1] hour resolution; size 9; (1,2] hour, size 7; (2,4] hour, size 5; and greater than 4, size 3. Endpoints outside the range are not included. For example, data at time point 4 with a 6-hour resolution would be compared to data at time points 3 and 5. This increasing window of comparison for smaller resolution identifies a true peak in the data, rather than simply noise. Endpoints are not included in peak calculations.

[1]Elements from the main text are referred to by M.X, where X is the element number.

The other support, the logarithmic decrement, is defined as the following:

$$\delta = \begin{cases} \delta = \frac{1}{n}\ln\frac{x(t)}{x(t+T)} & \text{if peak(1)} \ > \ \text{peak (2)} \\ \delta = \frac{1}{n}\ln\frac{x(t+nT)}{x(t)} & \text{otherwise} \end{cases} \tag{3}$$

where $n$ is the period between peaks, $t$ is the time of the first peak, and $t+T$ is the time of the next peak. The logarithmic decrement estimates the amount of damping between two peaks. Since we can think about forcing as reverse damping, we change the formulas for (1) and (3) accordingly.

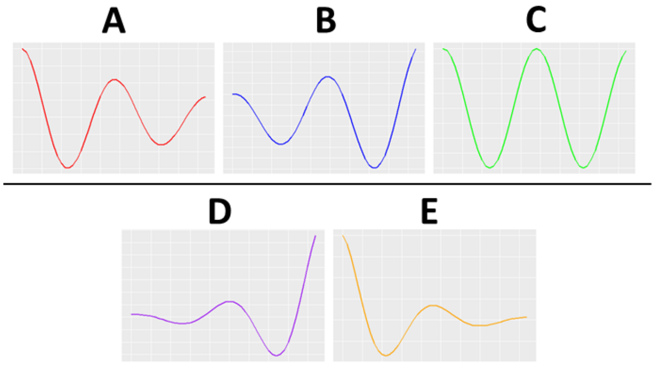


Figure 1: **Amplitude change coefficient determines differential categorizations of rhythms.** Representative curves for each of the AC coefficient categories. If $0.03 \leq \gamma \leq 0.15$, the rhythm is categorized as damped (A); $-0.15 \leq \gamma \leq -0.03$, the rhythm is categorized as forced (B); $-0.03 \leq \gamma \leq 0.03$, the rhythm is categorized as harmonic (C). In our analyses, these three categories are considered circadian. If $\gamma \leq -0.15$, the rhythm is categorized as overexpressed (D), and if $\gamma \geq 0.15$, the rhythm is categorized as repressed (E). In our analyses, the expression patterns in D and E are not considered circadian and are excluded.

#### 2 Representative Curves for Each AC Coefficient Category

Representative curves for each AC coefficient category appear in Fig. 1.

#### 3 Determining Cutoffs for Amplitude Change Coefficient Categories

We determined which ranges for $\gamma$, the amplitude change coefficient, were considered harmonic by examining several cutoffs for $\gamma$. When looking at representative curves for lower or higher values of the amplitude change coefficient, we considered the amount of amplitude loss for two cycles for each harmonic cutoff (Fig. 2A). We chose a conservative amount of greater than 50% amplitude loss, corresponding to an AC coefficient of $\pm.03$, in order to account for the large amounts of noise that can be seen in various biological datasets. However, it should be noted that this cutoff can be adjusted based on the amount of noise in the dataset. For datasets with less noise, this cutoff should be tightened, since a 'harmonic' rhythm corresponds to an ideal constant amplitude oscillator.

To determine cutoffs for non-circadian AC coefficient values, we followed a similar procedure by examining several cutoffs for $\gamma$. When looking at representative curves for lower or higher values of the amplitude change coefficient, we determined that anything greater than $\pm0.15$ could not be considered circadian for reasons cited in Section M2.1.4 (Fig. 2B). Further, we sought to confirm this in empirical observation using biological data from a highly sampled dataset from *Neurospora crassa* [10]. When we created heat maps of rhythms with a range of amplitude change coefficients, we noted that when the absolute value of the amplitude change coefficient was beyond $\pm0.15$, the expression patterns appeared less circadian than those with a amplitude change coefficient below 0.15 (Fig. 2C).

It should also be noted that these cutoffs may also be determined with confidence intervals; for example, if a confidence interval for the AC coefficient contained 0, it may be considered harmonic.

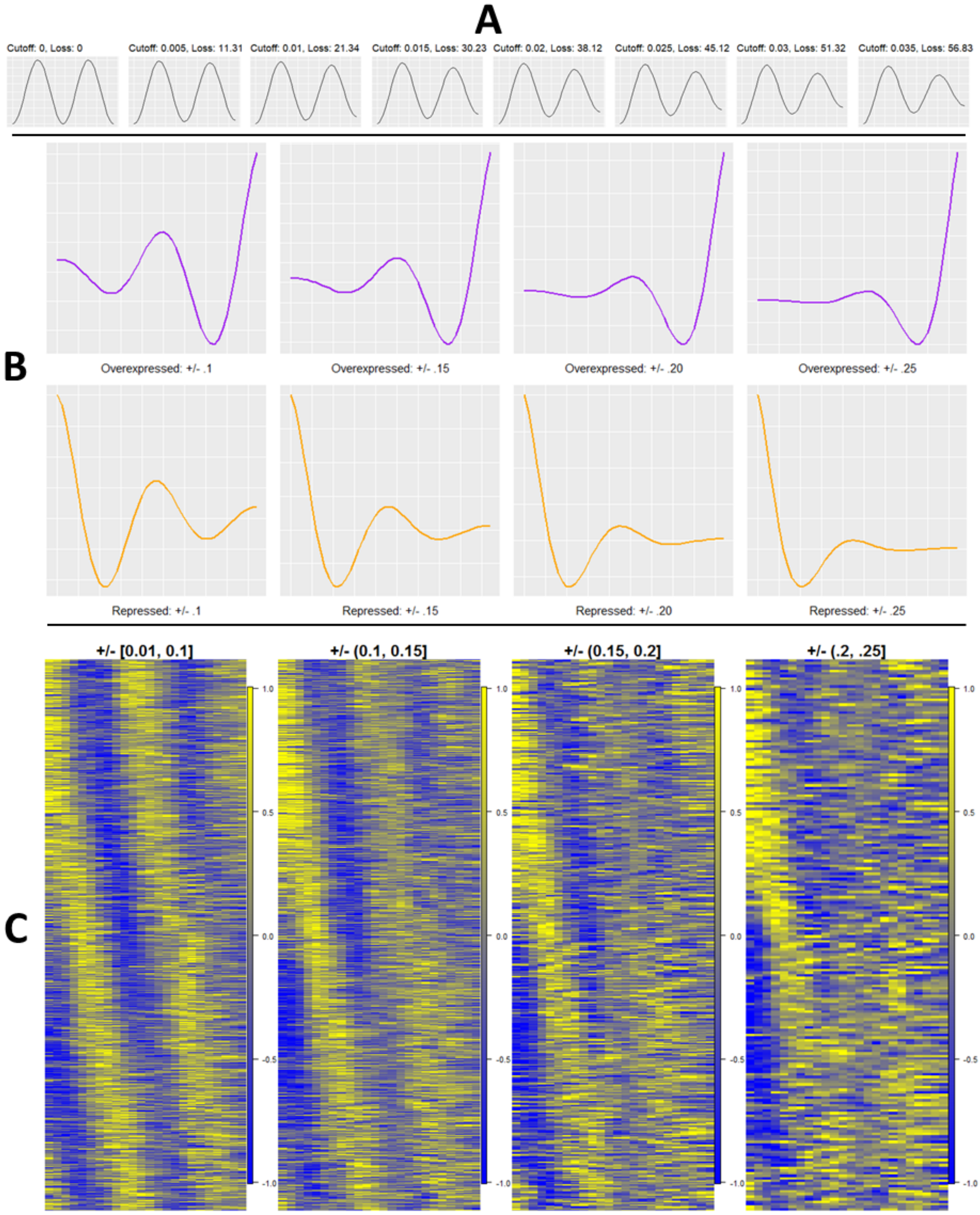


Figure 2: **Empirical analysis suggests amplitude change coefficient cutoffs.** A. Representative curves detailing the amplitude loss over two cycles for a sliding amplitude change coefficient cutoff, when considering the cutoff for harmonic rhythms. B. Representative curves for the specified threshold of amplitude change coefficient. The top row displays curves with positive amplitude change coefficients and the bottom row displays curves with negative amplitude change coefficients. C. Representative heat maps generated at each amplitude change coefficient level using data from *Neurospora crassa* transcripts [10]. Each row corresponds to the relative mean-centered expression of a single transcript and each column to a single time point. Heat map color keys are shown to the right of each map, which are bounded by -1 and 1, as each expression value is divided by the maximum magnitude of expression across the time course for each gene.

However, the exact implementation of the use of these confidence intervals remains unclear, as a confidence interval binning could also consider the ranges and cutoffs proposed above. Further, the calculation of confidence intervals through bootstrapping requires more computational time and does not provide consistency in small amounts of repetitions, since it is based on random sampling with replacement. Thus, we leave to future work the determination of AC categories by this method.

### 4 The Extended Circadian Harmonic Oscillator (ECHO) Application

#### 4.1 ECHO User Interface

In ECHO, under the first tab, "Find Rhythms", the user may upload data in .csv format, enter dataset information (including the time point range, resolution, and the amount and type of replicates), and can select to run JTK_CYCLE (JTK) in tandem with ECHO, as a comparison for current field standards (Fig. 3A) [9]. The user also has a choice of preprocessing steps to prepare their data for analysis (see below). As ECHO and JTK are run, a progress bar displays in the bottom right corner of the console window, and total run time is displayed at the top of the screen upon completion. The results can be downloaded as both .csv and .RData files. The .RData file also contains user inputs.

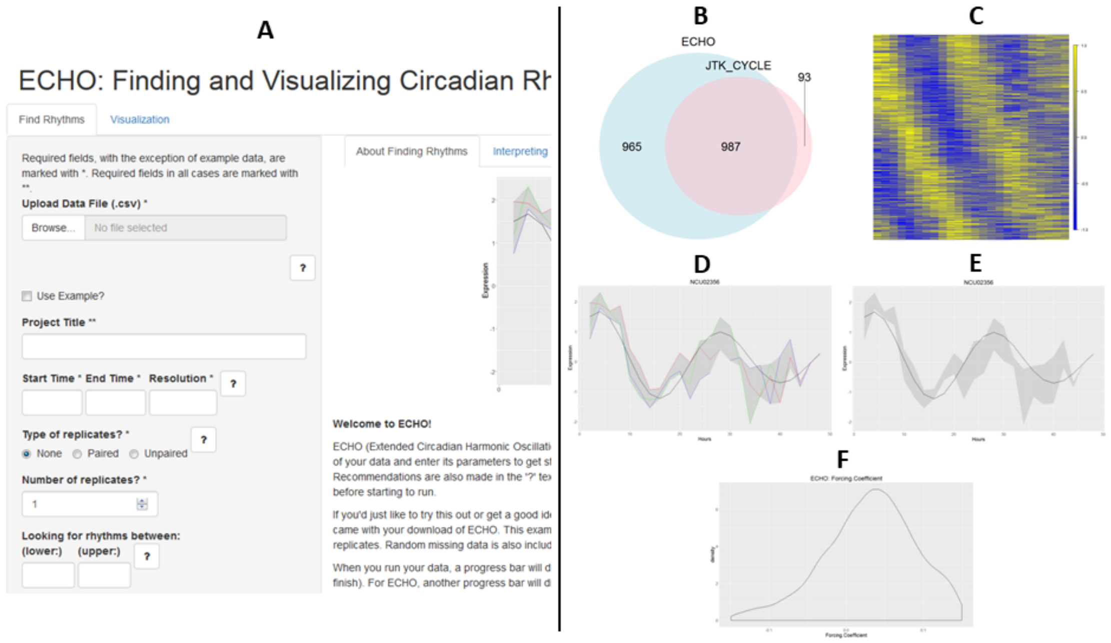


Figure 3: **The ECHO app provides a simple way for users to analyze and explore results.** A. The "Finding Rhythms" tab, where users enter dataset information and choose from a variety of preprocessing steps. B. A Venn Diagram produced by the ECHO app, for comparison between data from ECHO and JTK_CYCLE. C. Heat maps visualize average, mean-centered, relative expression among replicates, using yellow to indicate high expression and blue to indicate low expression. D. Gene expression plots display both the original and fitted data. For multiple replicates, the fit is displayed with shading between the maximum and the minimum values at that time point, with an option for the individual replicates to be displayed. E. Gene expression plot without each replicate displayed. F. Parameter density graphs display the distribution of a selected parameter.

The .RData file may then be uploaded in the second tab, "Visualizing Results", which allows for the exploration and visualization of the resultant data. The user can explore subsets of data, e.g. genes identified as rhythmic by ECHO. Moreover, the user can choose from a host of visualizations, including Venn diagrams, heat maps, gene expression plots, and parameter density graphs (Fig.

3B-F). ECHO allows for the specification of many visualization parameters, including significance levels, amplitude change coefficients, and period ranges. Images and gene lists can be exported for use in publication.

#### 4.2 Optional Preprocessing Steps in ECHO

Before fitting the data, ECHO has the option to perform several preprocessing steps. First, ECHO can smooth the data by implementing a running average over a 3-point window. This average is weighted at the center point of the window in the following manner:

$$x(t_0)_{smoothed} = \frac{2x(t_0) + x(t_1)}{3}$$
$$x(t_i)_{smoothed} = \frac{x(t_{i-1}) + 2x(t_i) + x(t_{i+1})}{4}$$
$$x(t_n)_{smoothed} = \frac{x(t_{n-1}) + 2x(t_n)}{3}$$

where $x(t_i)$ represents the expression at time $i = 0, ..., n$ and the denominator represents the sum of the weights. In using this weighted average, the data retains a larger proportion of the overall trend of the data than by using the standard running average, while at the same time retaining the noise-mitigating advantages of smoothing.

In addition to smoothing, ECHO can remove data with constant or undetected expression prior to analysis to save processing power. ECHO also has an option to remove genes that were detected in less than 70% of the experimental time points to eliminate experimental artifacts that can be caused by missing data [7]. Both options prevent negative effects on multiple hypothesis tested p-values.

To enable the direct comparison between data with vastly different amplitudes, ECHO provides a normalization option, which converts the expression at each time point to a z-score:

$$z_g(t_i) = \frac{x_g(t_i) - \mu_g}{\sigma_g} \tag{4}$$

where $\mu$ is the mean expression for a specific gene $g$, and $\sigma$ is the standard deviation of the expression for $g$. This creates a resultant $z$-score with a mean of 0 and a standard deviation of 1. In addition to enabling the comparison between genes, normalization reduces the size of the optimization space, reducing run time.

Finally, ECHO can remove linear trends from the data. This baseline removal is essential for strongly downward or upward sloping expressions, as our model (M.2) does not account for this and therefore could result in inaccurate parameter results. This is completed by linear least squares, where for each expression the squared difference between the fitted and experimental data is minimized. This results in an intercept ($a$) and slope ($b$), used in obtaining the detrended results in the following manner:

$$x(t_i)_{detrended} = x(t_i) - (a + bt_i) \text{ for } i = 1, ..., n$$

where $x(t)$ is the original value, $t$ is the time point, and $n$ is the total number of time points. For paired replicates, this linear fit is computed and removed from each replicate separately. For unpaired replicates, the linear trend is computed and removed from all replicates at once.

##### 4.2.1 Preprocessing Parameter Considerations

It should be noted that specific preprocessing schemes will affect the outcome of parameters, especially the amplitude change (AC) coefficient, among other parameters. Smoothing, while reducing noise, may attenuate the AC coefficient, as expressions at each time point will be dragged more towards the mean expression. Removing linear trends will change the AC coefficient in proportion with slope. For example, a rhythm with an initially strongly negative slope has a highly forced AC coefficient category as optimization attempts to minimize the squared residuals; once this slope is removed, the rhythm is damped instead. Normalization, however, does not change the AC coefficient. It only changes initial amplitude and equilibrium shift. This is displayed by the following:

$$
\begin{aligned}
x(t) &= \frac{(Ae^{\frac{-\gamma t}{2}}\cos(\omega t+\phi)+y)-\mu}{\sigma} \\
&= \frac{Ae^{\frac{-\gamma t}{2}}\cos(\omega t+\phi)+(y-\mu)}{\sigma} \\
&= \frac{A}{\sigma}e^{\frac{-\gamma t}{2}}\cos(\omega t+\phi)+\frac{y-\mu}{\sigma} \\
&= A_1 e^{\frac{-\gamma t}{2}}\cos(\omega t+\phi)+y_1
\end{aligned}
$$

where all parameters retain their meaning from (M.2) and (4), and $A_1$ and $y_1$ indicate updated values after normalization of initial amplitude and equilibrium shift, respectively.

#### 4.3 ECHO "Free Runs"

To explore rhythms beyond circadian, a user may leave the range of periods section blank, triggering a "free run". This free run allows ECHO to look for periods in the range of their time point resolution, with a minimum of 1 hour, to the length of their time course. This allows users to more easily see ultradian and infradian rhythms in addition to the standard circadian rhythms (Fig. 4A, B). Ultradian rhythms are shorter than a ~24 circadian rhythm, while infradian rhythms are longer.

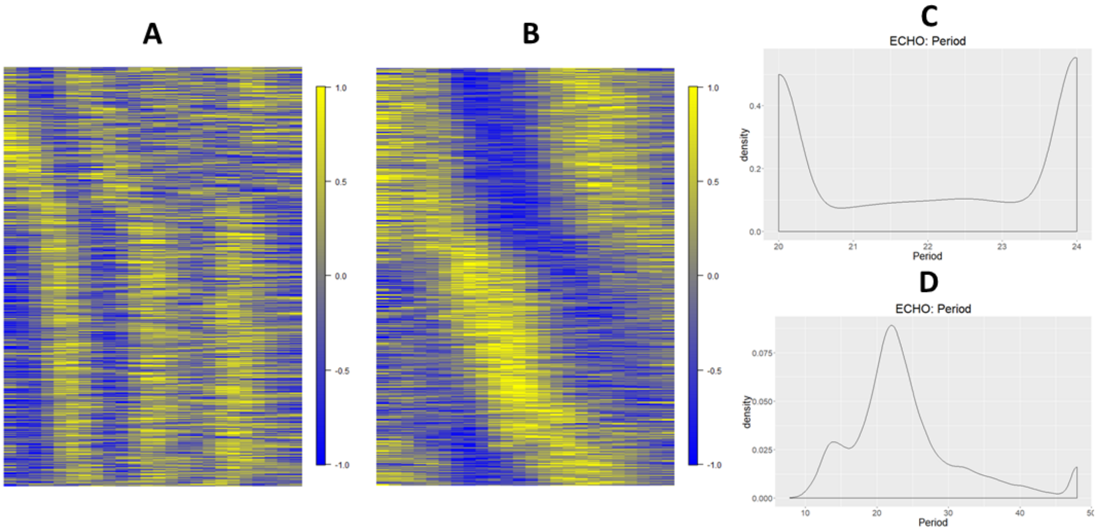


Figure 4: **ECHO "free runs" display optimal periods for ultradian and infradian rhythms in *Neurospora crassa*.** A, B. ECHO free run heatmaps of average, mean-centered, relative expression for mRNAs in the *Neurospora* transcriptome, processed as described in Section 4.2.1 [11]. Each row corresponds to the relative mean-centered expression of a single transcript and each column to a single time point. These heatmaps display an ultradian range of periods from 10 to 16 (A) and an infradian range of periods from 32 to 38 (B). C. Parameter density graph of period for a run of *Neurospora* where rhythms were specified to remain between 20 and 24 hours. D. Parameter density graph of period for a free run of *Neurospora*.

Further, a free run prevents ultradian or infradian genes that are passable at a certain p-value cutoff from being labeled circadian. In runs with ranges specified, parameter density graphs of the period often show high densities at either end of the range, indicating that these rhythms' more optimal periods likely lie outside this range, though this edge period provides a fit good enough to comply with the p-value cutoff (Fig. 4C). In a free run, a more optimal period can be identified for these rhythms (Fig. 4D).

### 5 Data

#### 5.1 Generation and Analysis of Synthetic Datasets

To evaluate ECHO's performance in recovering circadian genes as compared to other commonly used methods, we generated two sets of synthetic datasets using an approach similar to Wu et al and Deckard et al [16, 6]. Gene expression patterns were generated using (M.2) for each of our circadian AC categories. Initial parameters were randomly drawn from a uniform distribution with the following ranges:

$$A = \pm[8, 10]$$

$$\omega = \left[\frac{2\pi}{20}, \frac{2\pi}{30}\right]$$

$$\phi = [-2\pi, 2\pi]$$

$$y = [-4, 4]$$

The AC coefficient for each of the three categories (damped, forced, and harmonic) was also drawn from uniform distributions over their respective ranges. In addition, we added random noise to the data from normal distributions, centered at ranges categorized with respect to the amount of noise relative to the maximum possible initial amplitude. We chose a standard deviation of $\frac{1}{6}$ of these ranges, such that 99.7% of noise would fall between the given range. By categorizing noise in this manner, we provide a more intuitive method for interpreting noise. Low noise was drawn from a range of $[1, 2]$, medium from $[2, 3]$, and high from $[3, 5]$. These expression patterns were generated for a 2-hour resolution time course from 2 to 48 hours. Time points were then accordingly removed to create time courses with 4- and 6-hour resolutions. Non-periodic expression patterns were composed by flat, linear, and exponential curves with noise drawn from normal distributions.

The first datasets generated contained 10,000 expression patterns for each amplitude change coefficient, noise, and resolution category combination, with a 1:4 ratio of circadian to non-circadian genes, a higher total number of genes than in Wu et al. [16]. This higher total number of expressions is similar to what one might find in genome-wide studies, i.e., a large percentage of non-circadian genes with a smaller percentage of circadian genes and a larger number of genes total. The second datasets created used a less biologically relevant model by reducing the amount of noncircadian genes to 4,000 and increasing the ratio of circadian to non circadian genes to 1:1. However, this dataset allowed us to test the effects of multiple hypothesis testing on ECHO.

To analyze accuracy, we created ROC curves, plotting the true positive rate (TPR), or sensitivity, versus the false positive rate (FPR), or specificity. The Area Under the Curve (AUC) scores were calculated by subtracting the Benjamini-Hochberg adjusted p-values provided by each test from 1, such that a higher score would indicate a higher confidence in that expression being circadian. For MetaCycle, expressions' p-values were combined using Fisher's method, the suggested default (Supplemental Tables 1-6).

We then tested each program's accuracy in identifying the first peak in the data, or the phase. To do so, we computed the difference between the predicted phase shift by each program and the original generated phase shift for the simulated circadian genes that were detected by the program with a BH-adjusted p-value cutoff of 0.05. This data was arrayed using frequency polygon plots, which display the distribution across a histogram, binned for each hour. If the method is optimal, the distribution of each plot should be centered around 0, with sharply decreasing slopes on either side. We used a t-test to determine whether the mean distribution was significantly different from 0. Horizontal vertical lines at a magnitude of 10 indicate the range outside of which the difference corresponds to an antiphase prediction, as periods were originally drawn from $[20, 30]$ (Fig. 5-7).

#### 5.2 Acquisition and Pre-Processing of Biological Data Sets

Previously published data sets, including *Neurospora crassa* (RNAseq, SRP045821 and SRP046458; and Tandem Mass Tag proteomics, PXD009682 and 10.6019/PXD009682), Mouse liver (microarray)(GSE11923), NIH3t3 mouse embryo fibroblasts (microarray)(GSE11922), and *Anopheles gambiae* (microarray)(GSE22585) [10, 8, 12, 11] were acquired from publicly available databases. In the case of the *Neurospora* data sets, which did not include microarray data, the transcriptome and proteome were preprocessed as in Hurley et al. [10, 11], using the preprocessing steps described by Crowell et. al. [5]. Missing data in the transcriptomic dataset was imputed by averaging the time points on either side, in order to have the same structure as the completed proteomic data.

For the remaining microarray data sets, MAS5 and RMA normalization was completed using the Expression Console software from Affymetrix. The raw CEL files for each dataset were uploaded and run through both MAS5 and RMA following the instructions from the Expression Console, generating two output files per set. Each file was saved with an additional Gene ID column which became the main identifier as probe set IDs were removed. Present/absent calls from the MAS5 results and RMA normalization of the values were combined through an R script. Rows with more than 30% absent calls had all data points for those absent calls set to 0, while absent calls in rows with less than 30% remained at their original value.

In all data sets, we emulated the preprocessing steps in [8] and averaged all rows with the same identifying Gene ID. Data sets were analyzed using ECHO with parameters according to experimental design for sample resolution, number of replicates, and start and end time of the data collection. Transcripts were considered circadianly expressed if the Benjamini-Hochberg adjusted p-value was less than 0.05. AC coefficient ($\gamma$) cutoffs are as stated in Fig. 1. All datasets were analyzed with the following optional parameters within ECHO: data smoothing, unexpressed gene removal, data normalization, and a parallel run of JTK_CYCLE. For the animal data sets, we investigated a range of periods from 22-26 hours, while for the fungal datasets we used a 20-24-hour range, largely for compatibility with our comparison program, JTK_CYCLE [9]. However, ECHO program can be run as an unconstrained optimization problem; if no previous information is known about the period range of the oscillation, the program will search for rhythms as small as the resolution of the time course and as long as the time course length.

#### 5.3 Biological Analyses

##### 5.3.1 Gene Ontology (GO) Meta-Analysis

For each transcription-based dataset, damped, forced, and harmonic transcripts were determined via ECHO as previously described in Section 3.2 and evaluated for a statistical enrichment of gene ontologies using FungiFun2 (version 2.2.8) for *Neurospora crassa* transcripts, and Panther Classification System (version 14.0) for all other datasets using the appropriate reference organism and FDR-corrected p-values. For all terms enriched in at least one dataset, an average FDR-adjusted enrichment value for all datasets was calculated and multiplied by 2, as justified in [14].

##### 5.3.2 DREME Analysis for enriched motifs within Damped, Harmonic, and Forced oscillations in *Neurospora crassa*

To complete the DREME (Discriminative Regular Expression Motif Elicitation) analysis [1], we pulled promoter region sequences (here considered 500 bp before each transcription start site) for each subset of amplitudes, downloaded from FungiDB, Release 39 (30 Aug 2018) [3]. These 500 bp gene sets for forced, harmonic, and damped promoters, were each individually uploaded to the DREME program within Meme suite (meme-suite.org/) ver. 5.0.4, using shuffled input sequences as the control, the advanced option of searching only the given sequence and the default E-value cut-off of 0.05 [2, 1]. Discovered motifs were compared to a database of known *Neurospora crassa* transcription factors [15]. In all cases, plots for genes of interest were made using the ECHO app's visualization capabilities under its "Gene Expression with Replicates" option.

### 6 Results Auxiliary Figures and Tables

#### 6.1 Synthetic Data Results

AUCs for each method tested (ECHO, JTK_CYCLE, MetaCycle) appear in Tables 1 to 6. AUCs are truncated to 4 decimal places. Frequency polygon plots for differences between predicted and true phase appear in Fig. 5 to 7.

| | | Damped, 1:4 Ratio | | | | | | | | |
|---|---|---|---|---|---|---|---|---|---|---|
| | | Resolution | | | | | | | | |
| | | 2 hours | | | 4 hours | | | 6 hours | | |
| | | ECHO | JTK | MC | ECHO | JTK | MC | ECHO | JTK | MC |
| Noise | Low | 0.9234 | 0.9807 | 0.991 | 0.9063 | 0.8741 | 0.9236 | 0.8674 | 0.7005 | 0.8191 |
| | Medium | 0.8891 | 0.9146 | 0.9465 | 0.8627 | 0.7185 | 0.803 | 0.8163 | 0.5228 | 0.7099 |
| | High | 0.8357 | 0.7669 | 0.821 | 0.8082 | 0.5697 | 0.6714 | 0.7712 | 0.5041 | 0.631 |

Table 1: AUCs for synthetic data (S5.1) with a 1:4 ratio of damped circadian to noncircadian genes.

| | | Forced, 1:4 Ratio | | | | | | | | |
|---|---|---|---|---|---|---|---|---|---|---|
| | | Resolution | | | | | | | | |
| | | 2 hours | | | 4 hours | | | 6 hours | | |
| | | ECHO | JTK | MC | ECHO | JTK | MC | ECHO | JTK | MC |
| Noise | Low | 0.9883 | 1 | 0.9999 | 0.9754 | 0.998 | 0.9971 | 0.9313 | 0.8966 | 0.9539 |
| | Medium | 0.9868 | 1 | 1 | 0.9743 | 0.9976 | 0.9976 | 0.929 | 0.8934 | 0.9548 |
| | High | 0.982 | 1 | 0.9999 | 0.9722 | 0.997 | 0.9971 | 0.913 | 0.768 | 0.944 |

Table 2: AUCs for synthetic data (S5.1) with a 1:4 ratio of forced circadian to noncircadian genes.

| | | Harmonic, 1:4 Ratio | | | | | | | | |
|---|---|---|---|---|---|---|---|---|---|---|
| | | Resolution | | | | | | | | |
| | | 2 hours | | | 4 hours | | | 6 hours | | |
| | | ECHO | JTK | MC | ECHO | JTK | MC | ECHO | JTK | MC |
| Noise | Low | 0.9993 | 1 | 1 | 0.9971 | 0.9995 | 0.9998 | 0.9897 | 0.9617 | 0.9926 |
| | Medium | 0.999 | 0.9999 | 0.9999 | 0.9936 | 0.9939 | 0.9989 | 0.9794 | 0.9323 | 0.9804 |
| | High | 0.9902 | 0.9991 | 0.9999 | 0.9819 | 0.9426 | 0.9851 | 0.9417 | 0.7578 | 0.93 |

Table 3: AUCs for synthetic data (S5.1) with a 1:4 ratio split of harmonic circadian to noncircadian genes.

| | | Damped, 1:1 Ratio | | | | | | | | |
|---|---|---|---|---|---|---|---|---|---|---|
| | | Resolution | | | | | | | | |
| | | 2 hours | | | 4 hours | | | 6 hours | | |
| | | ECHO | JTK | MC | ECHO | JTK | MC | ECHO | JTK | MC |
| Noise | Low | 0.9138 | 0.9923 | 0.9968 | 0.9025 | 0.9243 | 0.9509 | 0.8541 | 0.8396 | 0.8576 |
| | Medium | 0.8859 | 0.9544 | 0.9682 | 0.8681 | 0.8377 | 0.8587 | 0.808 | 0.6978 | 0.7723 |
| | High | 0.839 | 0.8452 | 0.8567 | 0.8134 | 0.6938 | 0.7145 | 0.7702 | 0.589 | 0.6879 |

Table 4: AUCs for synthetic data (S5.1) with a 1:1 ratio of damped circadian to noncircadian genes.

| | | Forced, 1:1 Ratio | | | | | | | | |
|---|---|---|---|---|---|---|---|---|---|---|
| | | Resolution | | | | | | | | |
| | | 2 hours | | | 4 hours | | | 6 hours | | |
| | | ECHO | JTK | MC | ECHO | JTK | MC | ECHO | JTK | MC |
| Noise | Low | 0.9873 | 1 | 1 | 0.9804 | 0.9968 | 0.9972 | 0.9278 | 0.9579 | 0.962 |
| | Medium | 0.9876 | 1 | 1 | 0.979 | 0.9965 | 0.9962 | 0.9188 | 0.9514 | 0.9535 |
| | High | 0.9841 | 0.9999 | 0.9999 | 0.97 | 0.9956 | 0.9956 | 0.9207 | 0.9498 | 0.9557 |

Table 5: AUCs for synthetic data (S5.1) with a 1:1 ratio of forced circadian to noncircadian genes.

| | | Harmonic, 1:1 Ratio | | | | | | | | |
|---|---|---|---|---|---|---|---|---|---|---|
| | | **Resolution** | | | | | | | | |
| | | **2 hours** | | | **4 hours** | | | **6 hours** | | |
| | | **ECHO** | **JTK** | **MC** | **ECHO** | **JTK** | **MC** | **ECHO** | **JTK** | **MC** |
| **Noise** | **Low** | 0.9993 | 1 | 1 | 0.9978 | 0.9996 | 0.9999 | 0.9904 | 0.9793 | 0.9933 |
| | **Medium** | 0.9977 | 1 | 1 | 0.9955 | 0.9976 | 0.9988 | 0.9781 | 0.9625 | 0.9868 |
| | **High** | 0.9878 | 0.9994 | 0.9994 | 0.9779 | 0.9677 | 0.9864 | 0.939 | 0.9084 | 0.9496 |

Table 6: AUCs for synthetic data (S5.1) with a 1:1 ratio of harmonic circadian to noncircadian genes.

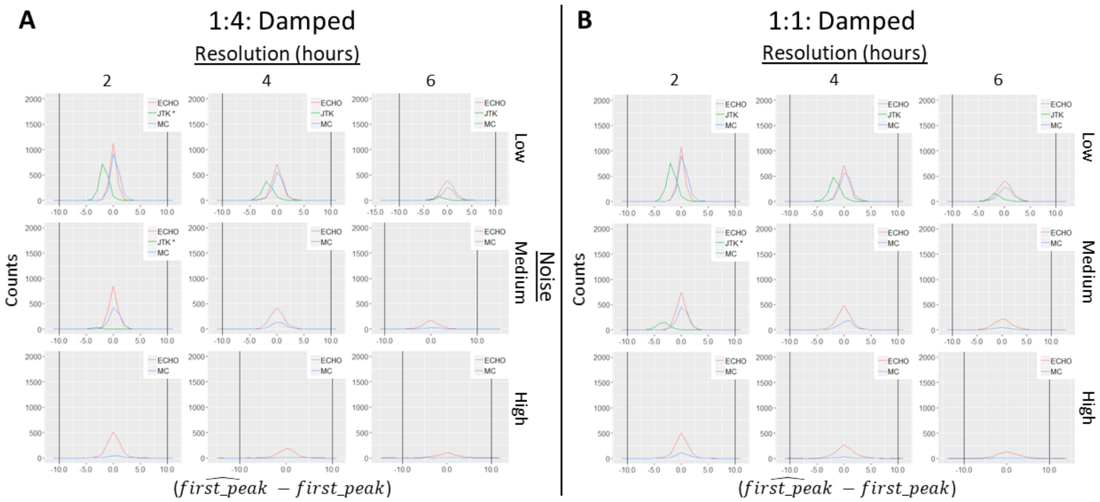


Figure 5: **Difference between predicted and actual phase frequency polygon plots for damped synthetic data.** Frequency polygon plots for damped synthetic data with a (A) 1:4 and a (B) 1:1 circadian to noncircadian ratio.

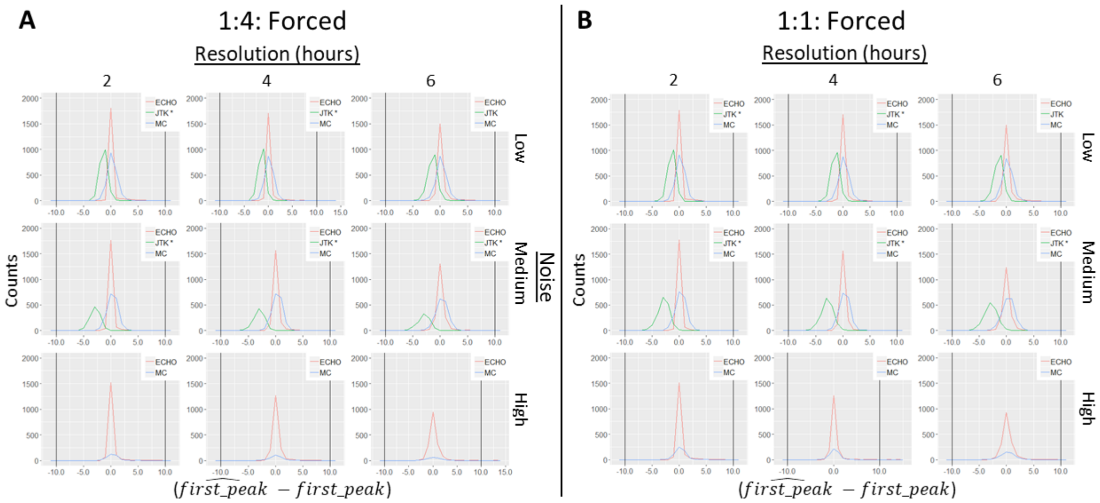


Figure 6: **Difference between predicted and actual phase frequency polygon plots for forced synthetic data.** Frequency polygon plots for forced synthetic data with a (A) 1:4 and a (B) 1:1 circadian to noncircadian ratio.

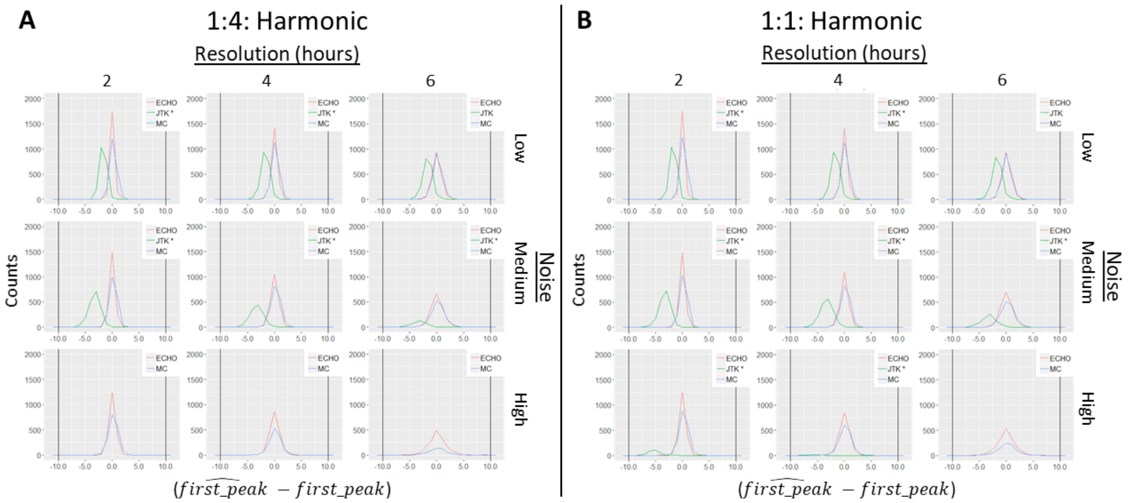


Figure 7: **Difference between predicted and actual phase frequency polygon plots for damped synthetic data.** Frequency polygon plots for harmonic synthetic data with a (A) 1:4 and a (B) 1:1 circadian to noncircadian ratio.

#### 6.2 Z-Score Visualization of Damped, Harmonic and Forced CCEs

To better visualize the overall expression profiles of damped, harmonic and forced CCEs identified by ECHO, we created heatmaps using z-scores of the expression value and plotted the average z-score at each time point for CCEs peaking in the same phase. The *Neurospora crassa* transcriptomic and *Anopheles gambiae* DD dataset are shown as representative examples in Fig. 8.

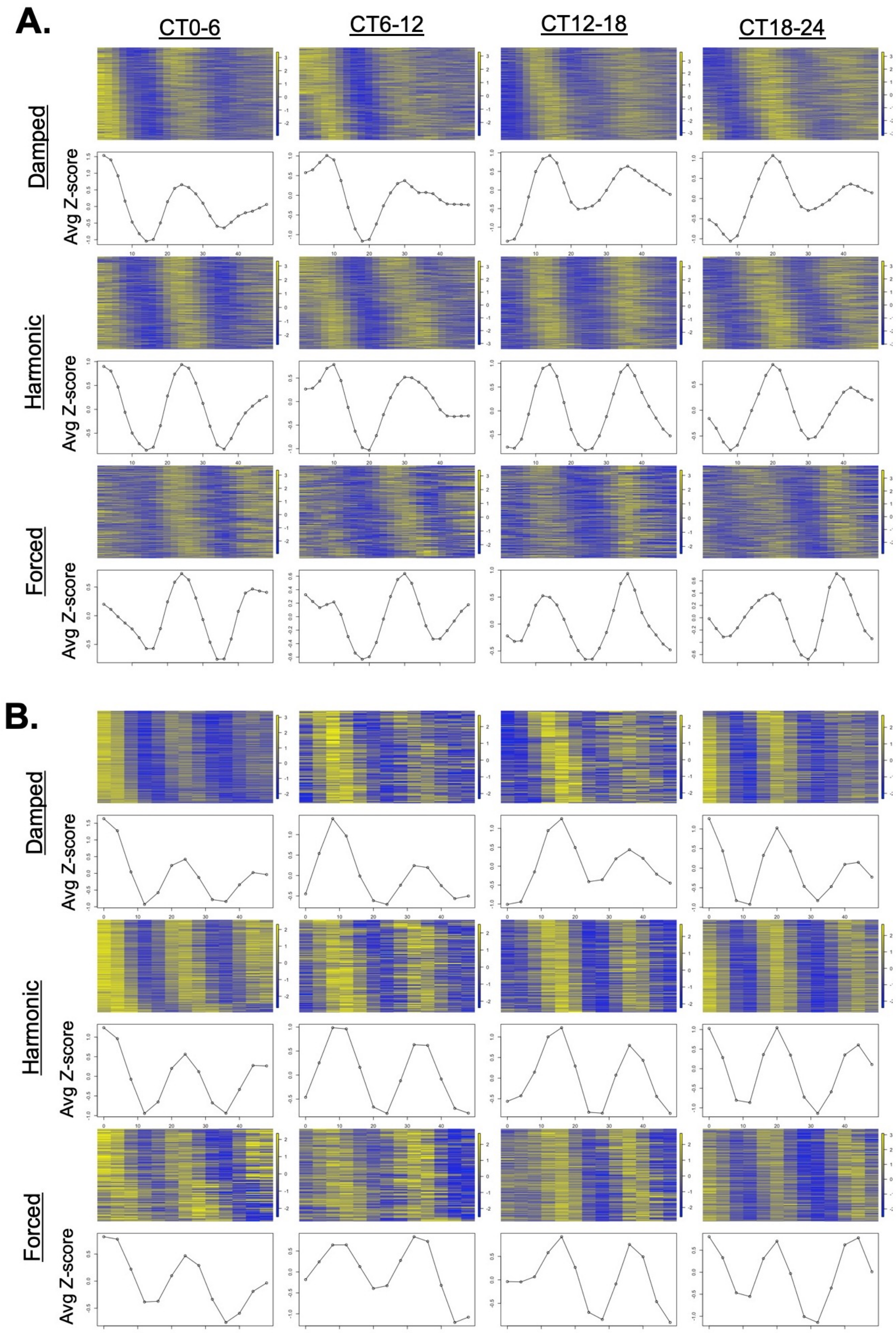


Figure 8: **Z-score heat maps and averages display clear AC coefficient rhythms.** Heat maps of different oscillation categories with z-score expression values for *Neurospora crassa* rhythmic transcripts (A) and *Anopheles gambiae* rhythmic transcripts in DD conditions (B), subset by circadian time of peak expression. Corresponding plots below heatmaps represent the average z-scores for all transcripts at each time point.

#### 6.3 Comparison of *Neurospora crassa* transcripts and proteins

A confusion matrix tabulating the amounts of rhythmic and non-rhythmic transcripts and proteins in *Neurospora crassa* appears in Table 7.

| | | Neurospora Protein | | | | | |
|---|---|---|---|---|---|---|---|
| | | **Damped** | **Forced** | **Harmonic** | **Overexpressed** | **Repressed** | **Not Rhythmic** |
| **Neurospora RNA** | **Damped** | 380 | 250 | 233 | 98 | 324 | 535 |
| | **Forced** | 82 | 52 | 69 | 33 | 71 | 165 |
| | **Harmonic** | 205 | 132 | 160 | 75 | 187 | 365 |
| | **Overexpressed** | 27 | 19 | 13 | 5 | 29 | 57 |
| | **Repressed** | 121 | 77 | 65 | 26 | 106 | 189 |
| | **Not Rhythmic** | 120 | 62 | 73 | 28 | 91 | 168 |

Table 7: A confusion matrix of *Neurospora crassa* transcripts and proteins, using data from [11].

#### 6.4 Comparison of Gene Ontologies for Damped, Forced, and Harmonic CCEs

All transcriptomic (microarray & RNA-seq derived) data described in Section M3.3 were included in a GO meta-analysis as described in Section 5.3.1. The top 20 ontologies based on P-value enrichment score unique to each category are listed in the Tables 8 to 10. All false discovery rates appearing the heat map in Fig. M.5 are given in Table SF.1[2].

| GO ID | PANTHER GO Term | Average FDR |
|---|---|---|
| GO:0005840 | ribosome | 5.30E-21 |
| GO:0005622 | intracellular | 4.90E-06 |
| GO:0019843 | rRNA binding | 1.01E-04 |
| GO:0006886 | intracellular protein transport | 9.66E-04 |
| GO:0006351 | transcription, DNA-templated | 1.12E-03 |
| GO:0004364 | glutathione transferase activity | 1.40E-03 |
| GO:0051179 | localization | 2.04E-03 |
| GO:0070925 | organelle assembly | 2.72E-03 |
| GO:0051128 | regulation of cellular component organization | 2.78E-03 |
| GO:0033036 | macromolecule localization | 3.28E-03 |
| GO:0003723 | RNA binding | 3.48E-03 |
| GO:0005886 | plasma membrane | 3.50E-03 |
| GO:0007010 | cytoskeleton organization | 4.34E-03 |
| GO:0016765 | transferase activity, transferring alkyl or aryl (other than methyl) groups | 4.40E-03 |
| GO:0016301 | kinase activity | 5.52E-03 |
| GO:0008104 | protein localization | 5.68E-03 |
| GO:0072593 | reactive oxygen species metabolic process | 8.64E-03 |
| GO:0034613 | cellular protein localization | 1.03E-02 |
| GO:0016773 | phosphotransferase activity, alcohol group as acceptor | 1.09E-02 |
| GO:0000226 | microtubule cytoskeleton organization | 1.12E-02 |

Table 8: **The top 20 unique gene ontological (GO) categories and their average false discovery rate for the damped rhythms subset.** A list of the top 20 unique GO terms retrieved from PANTHER for the damped subset, where the false discovery rate reported is an average false discovery rate from all datasets in which the term appeared, as described in Section 5.3.1.

[2]"SF" refers to supplemental file.

| GO ID | PANTHER GO Term | Average FDR |
|---|---|---|
| GO:0044444 | cytoplasmic part | 4.07E-04 |
| GO:0006397 | mRNA processing | 6.32E-04 |
| GO:0006412 | translation | 8.75E-04 |
| GO:0044451 | nucleoplasm part | 1.57E-03 |
| GO:0016043 | cellular component organization | 1.73E-03 |
| GO:0000180 | cytosolic large ribosomal subunit | 2.00E-03 |
| GO:0044391 | ribosomal subunit | 2.00E-03 |
| GO:0002181 | cytoplasmic translation | 2.19E-03 |
| GO:0003729 | mRNA binding | 3.60E-03 |
| GO:0006396 | RNA processing | 6.02E-03 |
| GO:0043161 | proteasome-mediated ubiquitin-dependent protein catabolic process | 6.45E-03 |
| GO:0016567 | protein ubiquitination | 1.01E-02 |
| GO:0016818 | hydrolase activity, acting on acid anhydrides, in phosphorus-containing anhydrides | 1.13E-02 |
| GO:0044445 | cytosolic part | 1.37E-02 |
| GO:0044446 | intracellular organelle part | 1.48E-02 |
| GO:0055085 | transmembrane transport | 1.49E-02 |
| GO:0016462 | pyrophosphatase activity | 1.56E-02 |
| GO:0005623 | cell | 1.57E-02 |
| GO:0044464 | cell part | 1.57E-02 |
| GO:0043632 | modification-dependent macromolecule catabolic process | 1.72E-02 |

Table 9: **The top 20 unique gene ontological (GO) categories and their average false discovery rate for the forced rhythms subset.** A list of the top 20 unique GO terms retrieved from PANTHER for the forced subset, where the false discovery rate reported is an average false discovery rate from all datasets in which the term appeared, as described in Section 5.3.1.

| GO ID | PANTHER GO Term | Average FDR |
|---|---|---|
| GO:0042254 | ribosome biogenesis | 9.28E-03 |
| GO:0030684 | preribosome | 9.66E-03 |
| GO:0043170 | macromolecule metabolic process | 1.56E-02 |
| GO:0015078 | proton transmembrane transporter activity | 1.71E-02 |
| GO:0016788 | hydrolase activity, acting on ester bonds | 1.75E-02 |
| GO:0015036 | disulfide oxidoreductase activity | 2.74E-02 |
| GO:0003779 | actin binding | 2.74E-02 |
| GO:0034220 | ion transmembrane transport | 2.76E-02 |
| GO:0004866 | endopeptidase inhibitor activity | 2.82E-02 |
| GO:0061135 | endopeptidase regulator activity | 2.94E-02 |
| GO:0061134 | peptidase regulator activity | 3.06E-02 |
| GO:0034455 | t-UTP complex | 3.20E-02 |
| GO:0005856 | cytoskeleton | 3.20E-02 |
| GO:0015629 | actin cytoskeleton | 3.26E-02 |
| GO:0005215 | transporter activity | 3.30E-02 |
| GO:0002020 | protease binding | 3.42E-02 |
| GO:0006364 | rRNA processing | 3.70E-02 |
| GO:0006511 | ubiquitin-dependent protein catabolic process | 3.74E-02 |
| GO:0030663 | COPI-coated vesicle membrane | 3.74E-02 |
| GO:0031410 | cytoplasmic vesicle | 3.74E-02 |

Table 10: **The top 20 unique gene ontological (GO) categories and their average false discovery rate for the harmonic rhythms subset.** A list of the top 20 unique GO terms retrieved from PANTHER for the harmonic subset, where the false discovery rate reported is an average false discovery rate from all datasets in which the term appeared, as described in Section 5.3.1.

#### 6.5 Damped and Forced Oscillations in *Neurospora crassa*

Lighting cues are also important in setting the phase of the molecular clock in *Neurospora* [4]. Since rhythms are known to dampen in some cases without continued external cues, such as light [7], some of the damped rhythms in *Neurospora* could be related to the absence of lighting cues. Indeed the core clock genes *Neurospora wc-1* and *wc-2*, were classified in ECHO as repressed or damped, respectively (Fig. 9). Previously identified WCC targets [13, 10] were also mostly damped or repressed, suggesting that the damping rhythm from the WCC is being propagated to its target genes.

Using DREME analysis [1], a number of motifs were discovered within each ECHO-identified *Neurospora crassa* transcriptomic gene set: forced, n= 1069 genes, 34 motifs; harmonic, n= 2188 genes, 48 motifs; and damped, n= 3428 genes, 62 motifs. Targets of CRE-1, discussed in the main paper, appear here in Fig. 10.

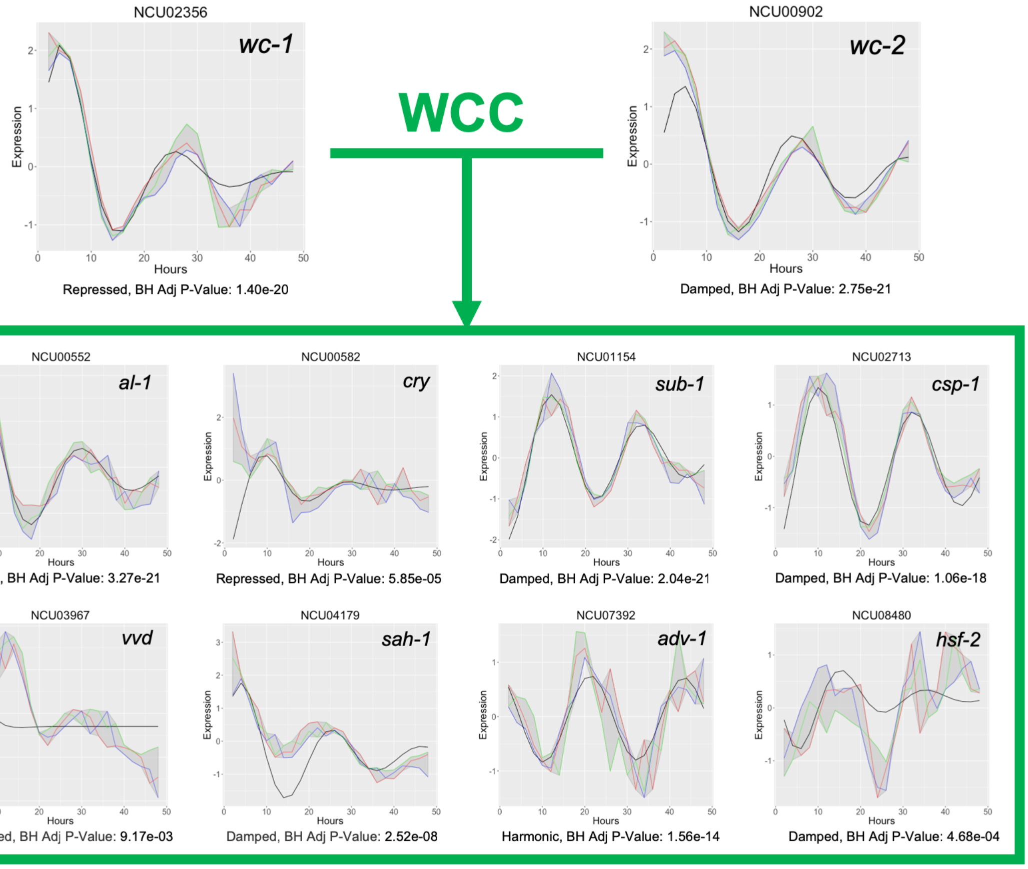


Figure 9: **Damping of the positive arm of the *Neurospora* clock (WCC) and its targets may be related to the damping of light induced pathways.** ECHO app-generated plots of *Neurospora* RNA-seq data (colored lines) and ECHO best-fit models (black line) for *wc-1* and *wc-2* and a select group of the genes they transcriptionally activate upon light induction (in green box). Gene IDs and gene names can be found at the top of each plot, and the best-fit model (harmonic, damped, or forced) and BH-adjusted P-values from ECHO are reported below each plot.

### 7 Software Version Notes

All data was run with ECHO interface v3.1, JTK_CYCLE v3.1, and MetaCycle v1.1.0, unless otherwise noted. Version updates and specifications for ECHO can be found on GitHub[3].

[3] https://github.com/delosh653/ECHO

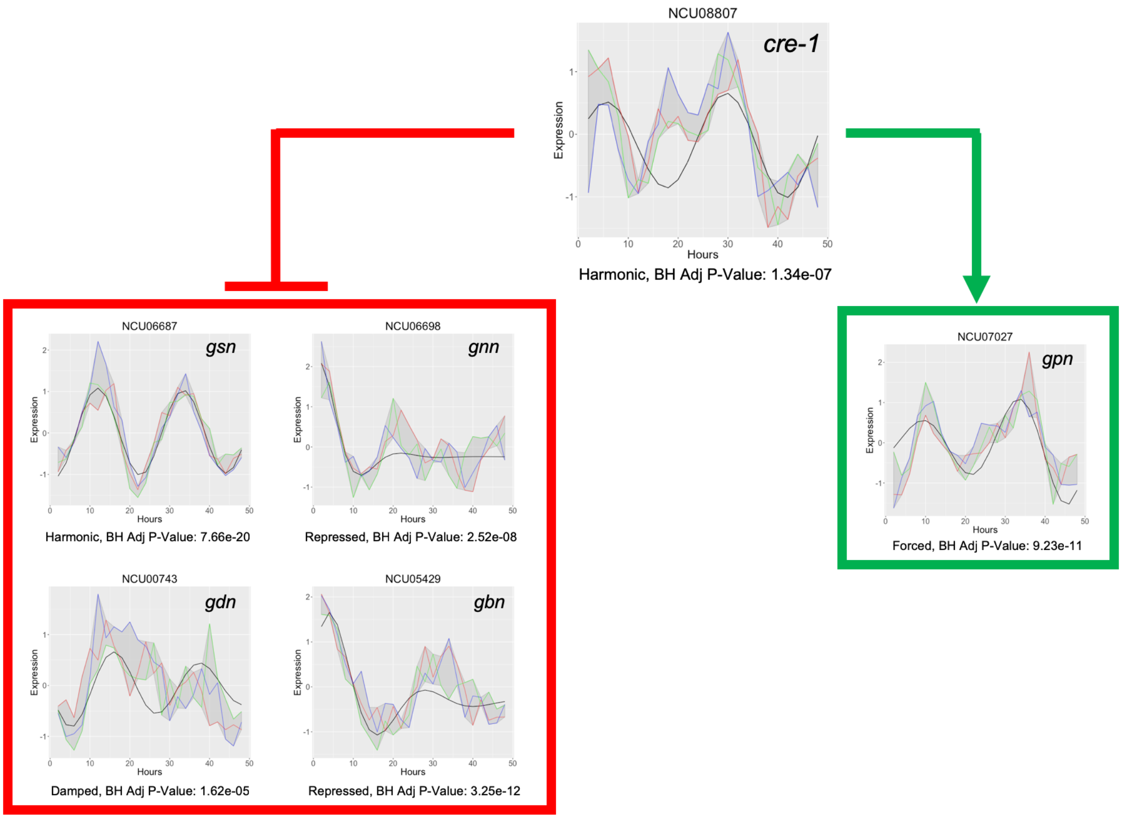


Figure 10: **ECHO identifies regulatory relationship between *cre-1* and its known targets.** ECHO app-generated plots of *Neurospora* RNA-seq data (colored lines) and ECHO best-fit models (black line) for *cre-1* and a select group of the genes it transcriptionally represses (in red box) or activates (in green box). Gene IDs and gene names can be found at the top of each plot, and the best-fit model (harmonic, damped, or forced) and BH-adjusted P-values from ECHO are reported below each plot.

[4] Susan K Crosthwaite, Jennifer J Loros, and Jay C Dunlap. Light-induced resetting of a circadian clock is mediated by a rapid increase in frequency transcript. *Cell*, 81(7):1003–1012, jun 1995.

[5] Alexander M Crowell, Jay C Dunlap, Casey S Greene, and Jennifer J Loros. Learning and Imputation for Mass-spec Bias Reduction (LIMBR). *Bioinformatics*, 09 2018.

[6] Anastasia Deckard, John Harer, John B. Hogenesch, Ron C. Anafi, and Steven B. Haase. Design and analysis of large-scale biological rhythm studies: a comparison of algorithms for detecting periodic signals in biological data. *Bioinformatics*, 29(24):3174–3180, 09 2013.

[7] Michael E Hughes, Katherine C Abruzzi, Ravi Allada, Ron Anafi, Alaaddin Bulak Arpat, Gad Asher, Pierre Baldi, Charissa de Bekker, Deborah Bell-Pedersen, Justin Blau, Steve Brown, M Fernanda Ceriani, Zheng Chen, Joanna C Chiu, Juergen Cox, Alexander M Crowell, Jason P DeBruyne, Derk-Jan Dijk, Luciano DiTacchio, Francis J Doyle, Giles E Duffield, Jay C Dunlap, Kristin Eckel-Mahan, Karyn A Esser, Garret A FitzGerald, Daniel B Forger, Lauren J Francey, Ying-Hui Fu, Frédéric Gachon, David Gatfield, Paul de Goede, Susan S Golden, Carla Green, John Harer, Stacey Harmer, Jeff Haspel, Michael H Hastings, Hanspeter Herzel, Erik D Herzog, Christy Hoffmann, Christian Hong, Jacob J Hughey, Jennifer M Hurley, Horacio O de la Iglesia, Carl Johnson, Steve A Kay, Nobuya Koike, Karl Kornacker, Achim Kramer, Katja Lamia, Tanya Leise, Scott A Lewis, Jiajia Li, Xiaodong Li, Andrew C Liu, Jennifer J Loros, Tami A Martino, Jerome S Menet, Martha Merrow, Andrew J Millar, Todd Mockler, Felix Naef, Emi Nagoshi, Michael N Nitabach, Maria Olmedo, Dmitri A Nusinow, Louis J Ptáček, David Rand, Akhilesh B Reddy, Maria S Robles, Till Roenneberg, Michael Rosbash, Marc D Ruben, Samuel S C Rund, Aziz Sancar, Paolo Sassone-Corsi, Amita Sehgal,

Scott Sherrill-Mix, Debra J Skene, Kai-Florian Storch, Joseph S Takahashi, Hiroki R Ueda, Han Wang, Charles Weitz, Pål O Westermark, Herman Wijnen, Ying Xu, Gang Wu, Seung-Hee Yoo, Michael Young, Eric Erquan Zhang, Tomasz Zielinski, and John B Hogenesch. Guidelines for Genome-Scale Analysis of Biological Rhythms. *Journal of Biological Rhythms*, 32(5):380–393, oct 2017.

[8] Michael E. Hughes, Luciano DiTacchio, Kevin R. Hayes, Christopher Vollmers, S. Pulivarthy, Julie E. Baggs, Satchidananda Panda, and John B. Hogenesch. Harmonics of circadian gene transcription in mammals. *PLoS Genetics*, 2009.

[9] Michael E. Hughes, John B. Hogenesch, and Karl Kornacker. JTK-CYCLE: An efficient non-parametric algorithm for detecting rhythmic components in genome-scale data sets. *Journal of Biological Rhythms*, 2010.

[10] Jennifer M Hurley, Arko Dasgupta, Jillian M Emerson, Xiaoying Zhou, Carol S Ringelberg, Nicole Knabe, Anna M Lipzen, Erika A Lindquist, Christopher G Daum, Kerrie W Barry, Igor V Grigoriev, Kristina M Smith, James E Galagan, Deborah Bell-Pedersen, Michael Freitag, Chao Cheng, Jennifer J Loros, and Jay C Dunlap. Analysis of clock-regulated genes in Neurospora reveals widespread posttranscriptional control of metabolic potential. *Proceedings of the National Academy of Sciences of the United States of America*, 2014.

[11] Jennifer M. Hurley, Meaghan S. Jankowski, Hannah De los Santos, Alexander Crowell, Sam Fordyce, Jeremy D. Zucker, Neeraj Kumar, Sam Purvine, Errol Robinson, Anil Shukla, Erika Zink, William R. Cannon, Scott E. Baker, Jennifer J. Loros, and Jay C. Dunlap. Circadian proteomic analysis uncoversmechanisms of post-transcriptionalregulation in metabolic pathways. *Cell Systems*, 2018.

[12] S. S. C. Rund, T. Y. Hou, S. M. Ward, F. H. Collins, and G. E. Duffield. Genome-wide profiling of diel and circadian gene expression in the malaria vector Anopheles gambiae. *Proceedings of the National Academy of Sciences*, 108(32):E421–E430, 2011.

[13] Kristina M. Smith, Gencer Sancar, Rigzin Dekhang, Christopher M. Sullivan, Shaojie Li, Andrew G. Tag, Cigdem Sancar, Erin L. Bredeweg, Henry D. Priest, Ryan F. McCormick, Terry L. Thomas, James C. Carrington, Jason E. Stajich, Deborah Bell-Pedersen, Michael Brunner, and Michael Freitag. Transcription factors in light and circadian clock signaling networks revealed by genomewide mapping of direct targets for neurospora white collar complex. *Eukaryotic Cell*, 9(10):1549–1556, 2010.

[14] Vladimir Vovk and Ruodu Wang. Combining P-Values Via Averaging. *SSRN Electronic Journal*, 2018.

[15] Matthew T. Weirauch, Ally Yang, Mihai Albu, Atina G. Cote, Alejandro Montenegro-Montero, Philipp Drewe, Hamed S. Najafabadi, Samuel A. Lambert, Ishminder Mann, Kate Cook, and et al. Determination and inference of eukaryotic transcription factor sequence specificity. *Cell*, 158(6):1431–1443, 2014.

[16] Gang Wu, Jiang Zhu, Jun Yu, Lan Zhou, Jianhua Z. Huang, and Zhang Zhang. Evaluation of five methods for genome-wide circadian gene identification. *Journal of Biological Rhythms*, 29(4):231–242, 2014.
